# Perturb-seq identifies TCF7 as a central nexus linking MAPK- and Wnt-driven gene expression

**DOI:** 10.64898/2025.12.15.694273

**Authors:** Ghanem El Kassem, Anja Sieber, Bertram Klinger, Florian Uhlitz, David Steinbrecht, Mirjam van Bentum, Jasmine Hillmer, Jennifer von Schlichting, Reinhold Schäfer, Nils Blüthgen, Michael Boettcher

## Abstract

The MAPK pathway is a central signaling cascade whose dysregulation contributes to numerous diseases. While its upstream regulation is well studied, the mechanisms by which MAPK activation leads to diverse transcriptional outcomes remain incompletely understood. To address this shortcoming, we mapped the target gene sets controlled by 22 RAF-inducible transcription factors using targeted Perturb-seq and integrated these data with time-resolved transcriptional profiling. Network reconstruction revealed a topology dominated by two central hubs, EGR1 and FOS, which co-regulate partially overlapping target gene sets. In addition, we uncovered a positive feedback loop between EGR1, a canonical RAF-MAPK effector, and TCF7, a transcription factor typically linked to Wnt signaling. Through this interaction, TCF7 emerges as a nexus that integrates MAPK and Wnt pathway inputs. Together, these findings define the architecture of RAF-MAPK-driven transcriptional regulation and demonstrate how cross-talk between oncogenic signaling pathways can be encoded in transcriptional networks.

**GRAPHICAL ABSTRACT:** 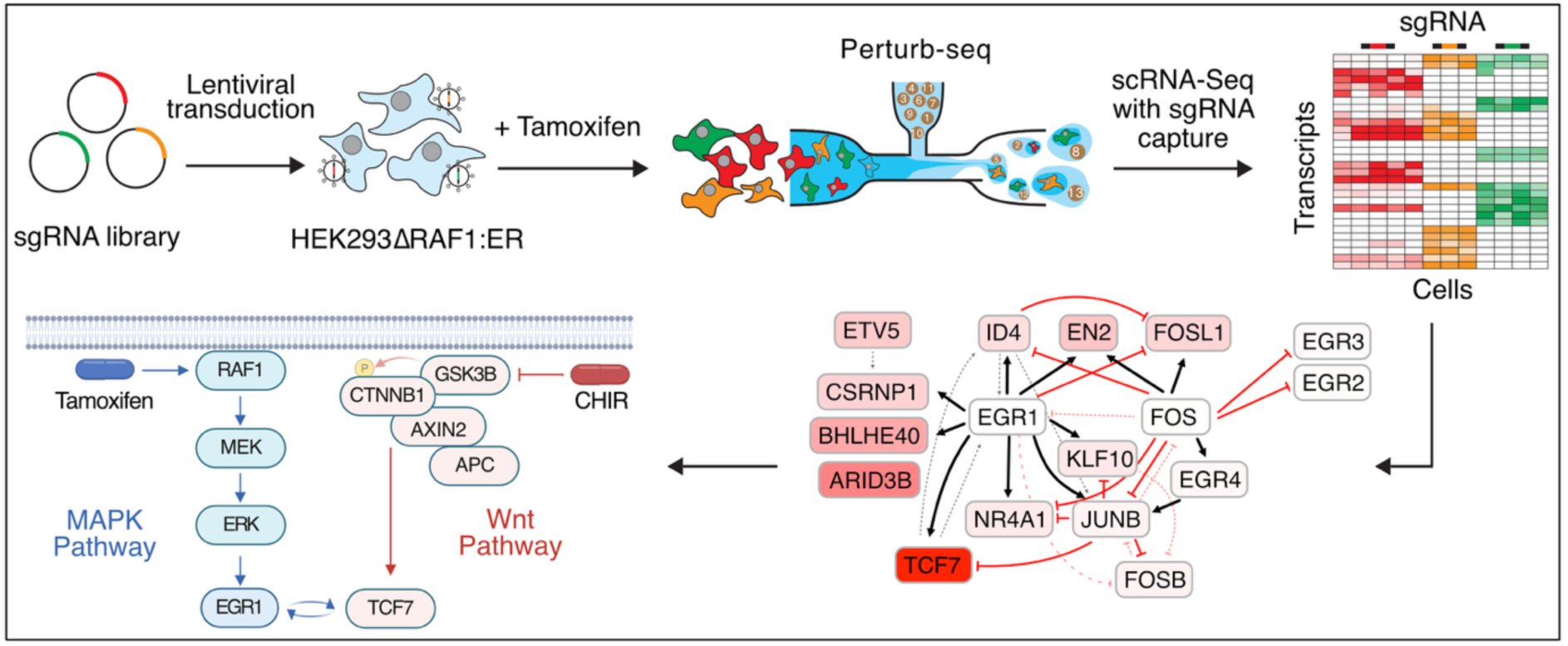

## INTRODUCTION

The Mitogen-Activated Protein Kinase (MAPK) signaling cascade is a fundamental and evolutionarily conserved pathway that regulates proliferation, differentiation, and survival (Cargnello & Roux, 2011). Activated by diverse external cues - including growth factors, hormones, and stress - MAPK transmits signals from the cell surface to the nucleus, where MAPKs phosphorylate transcription factors and initiate defined transcriptional programs. This adaptability allows cells to translate a broad spectrum of stimuli into context-specific responses.

The MAPK pathway has long served as a paradigm for studying signal-induced transcriptional programs (1). Early transcriptome studies after stimulation of the pathway led to the concepts of immediate-early and delayed primary genes that are activated by pre-existing transcription factors (2), and share regulatory motifs in their promoters (3, 4). Many immediate-early genes are transcription factors, which then induce secondary targets. Interestingly, delayed primary genes often encode negative feedback regulators (5–7), and the timing of primary response genes is strongly determined by mRNA half-lives (8). While the kinetics of the transcriptome response to MAPK activation has been characterized in quantitative depth (2, 5, 7, 8), the wiring of how immediate-early transcription factors induce secondary response genes and the understanding of their interaction remain cryptic. A better understanding of the topology of those transcriptional networks will be crucial for understanding the coordinated regulation of oncogenic pathways, with potential implications for therapeutic intervention.

Aberrant MAPK signaling is a hallmark of many diseases, particularly cancer (9, 10). For example, driver EGFR gene mutations occur in roughly 10-35% of non-small cell lung cancers, most frequently as exon 19 deletions or the L858R substitution in exon 21 (11). Activating mutations in KRAS that lead to aberrant activation of the MAPK signaling pathway are the most common with approximately 30% of all cancers carrying driver Ras mutations (12). These are followed by B-RAF mutations which account for around 8% of cancer cases, especially in melanoma and thyroid cancers (11). Reverse engineering of the transcriptional networks downstream of MAPK signaling represents a promising approach to better understand how physiological and pathological MAPK signals are integrated into a cellular response. We have previously used systematic perturbation data and reverse engineering to elucidate a small seven node transcription factor network downstream of RAS/MAPK signaling that controls transformation and different aspects of cell growth (13). Expansion of similar approaches to larger networks has been hampered by technical challenges for a long time. However, newly developed methods that combine CRISPR-based genetic perturbation techniques with single-cell RNA-Seq readout, such as Perturb-seq (14, 15), now enable simultaneous functional genetic perturbation and investigation of the resulting transcriptional response at scale.

Here, we systematically analyzed 22 RAF/MAPK-inducible transcription factors via targeted Perturb-seq (TAP-seq; (16)) to map the transcriptional programs downstream of the MAPK effector kinase RAF1. We integrated the results with time-resolved expression data to reconstruct the network topology. This analysis revealed two central hubs, EGR1 and FOS, which co-regulate overlapping and partially orthogonal target modules. Most notably, we discovered a positive feedback loop between EGR1 and TCF7, a transcription factor classically linked to Wnt signaling, despite their distinct induction kinetics. Pathway-level analysis demonstrated that this interaction enhances Wnt output, establishing TCF7 as a nexus of MAPK–Wnt cross-talk. Together, these findings delineate the architecture of RAF-MAPK-driven transcriptional programs and reveal a transcriptional mechanism that integrates two major oncogenic pathways.

## MATERIAL AND METHODS

### Reagents

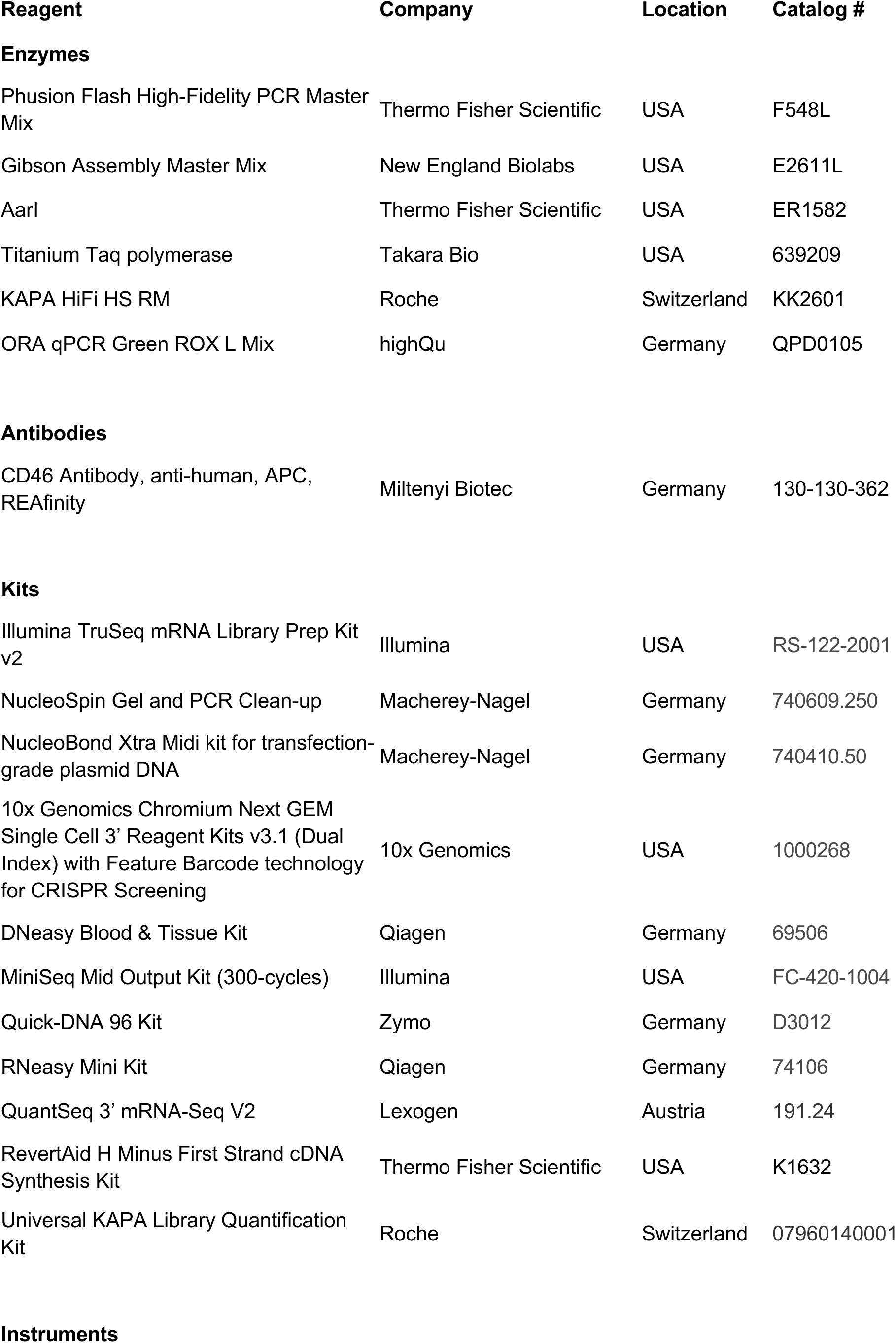
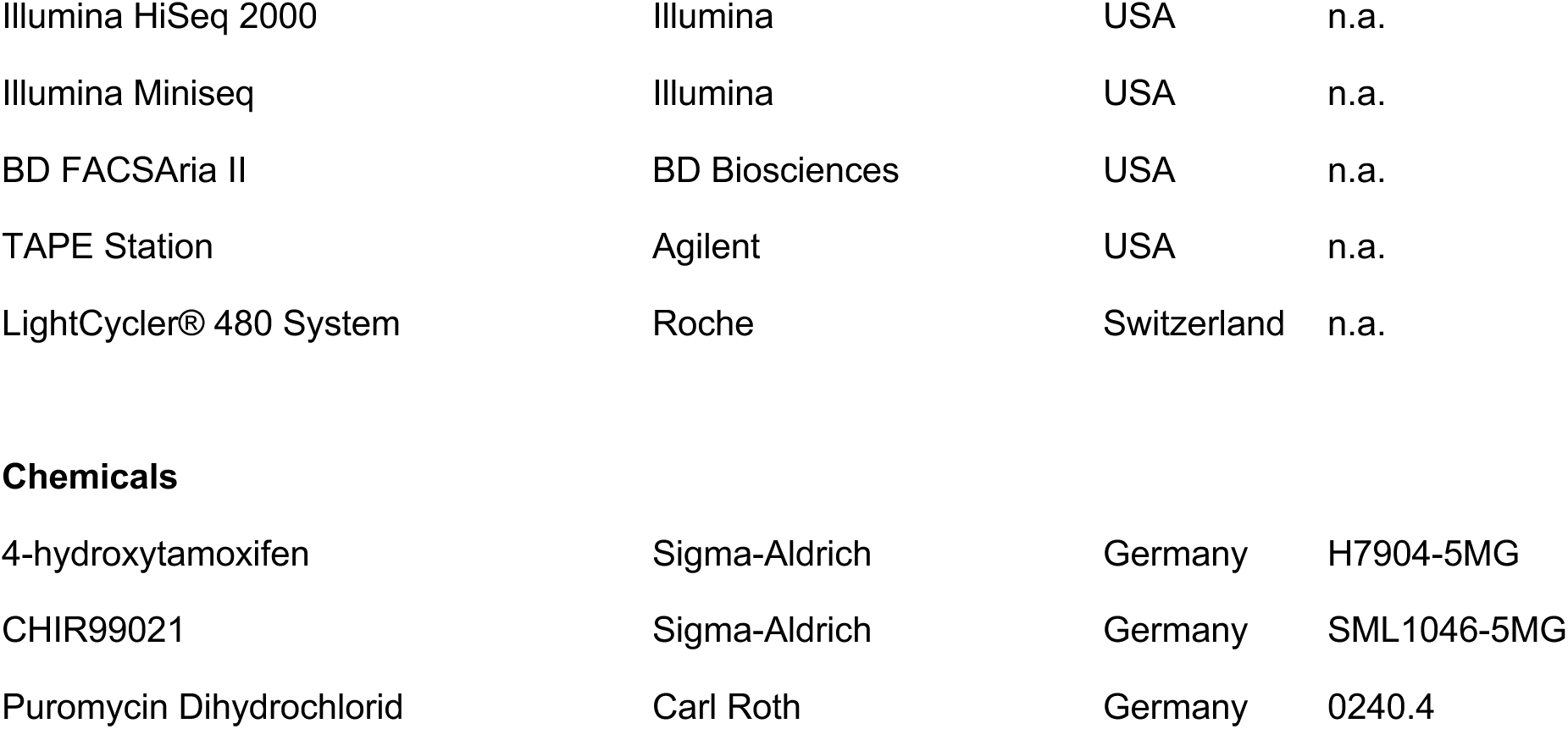

| Reagent | Company | Location | Catalog # |
| --- | --- | --- | --- |
| **Enzymes** |  |  |  |
| Phusion Flash High-Fidelity PCR Master Mix | Thermo Fisher Scientific | USA | F548L |
| Gibson Assembly Master Mix | New England Biolabs | USA | E2611L |
| AarI | Thermo Fisher Scientific | USA | ER1582 |
| Titanium Taq polymerase | Takara Bio | USA | 639209 |
| KAPA HiFi HS RM | Roche | Switzerland | KK2601 |
| ORA qPCR Green ROX L Mix | highQu | Germany | QPD0105 |
| **Antibodies** |  |  |  |
| CD46 Antibody, anti-human, APC, REAfinity | Miltenyi Biotec | Germany | 130-130-362 |
| **Kits** |  |  |  |
| Illumina TruSeq mRNA Library Prep Kit v2 | Illumina | USA | RS-122-2001 |
| NucleoSpin Gel and PCR Clean-up | Macherey-Nagel | Germany | 740609.250 |
| NucleoBond Xtra Midi kit for transfection-grade plasmid DNA | Macherey-Nagel | Germany | 740410.50 |
| 10x Genomics Chromium Next GEM Single Cell 3' Reagent Kits v3.1 (Dual Index) with Feature Barcode technology for CRISPR Screening | 10x Genomics | USA | 1000268 |
| DNeasy Blood & Tissue Kit | Qiagen | Germany | 69506 |
| MiniSeq Mid Output Kit (300-cycles) | Illumina | USA | FC-420-1004 |
| Quick-DNA 96 Kit | Zymo | Germany | D3012 |
| RNeasy Mini Kit | Qiagen | Germany | 74106 |
| QuantSeq 3' mRNA-Seq V2 | Lexogen | Austria | 191.24 |
| RevertAid H Minus First Strand cDNA Synthesis Kit | Thermo Fisher Scientific | USA | K1632 |
| Universal KAPA Library Quantification Kit | Roche | Switzerland | 07960140001 |
| Instruments |  |  |  |
| Illumina HiSeq 2000 | Illumina | USA | n.a. |
| Illumina Miniseq | Illumina | USA | n.a. |
| BD FACSAria II | BD Biosciences | USA | n.a. |
| TAPE Station | Agilent | USA | n.a. |
| LightCycler® 480 System | Roche | Switzerland | n.a. |
| Chemicals |  |  |  |
| 4-hydroxytamoxifen | Sigma-Aldrich | Germany | H7904-5MG |
| CHIR99021 | Sigma-Aldrich | Germany | SML1046-5MG |
| Puromycin Dihydrochlorid | Carl Roth | Germany | 0240.4 |

### Biological Resources

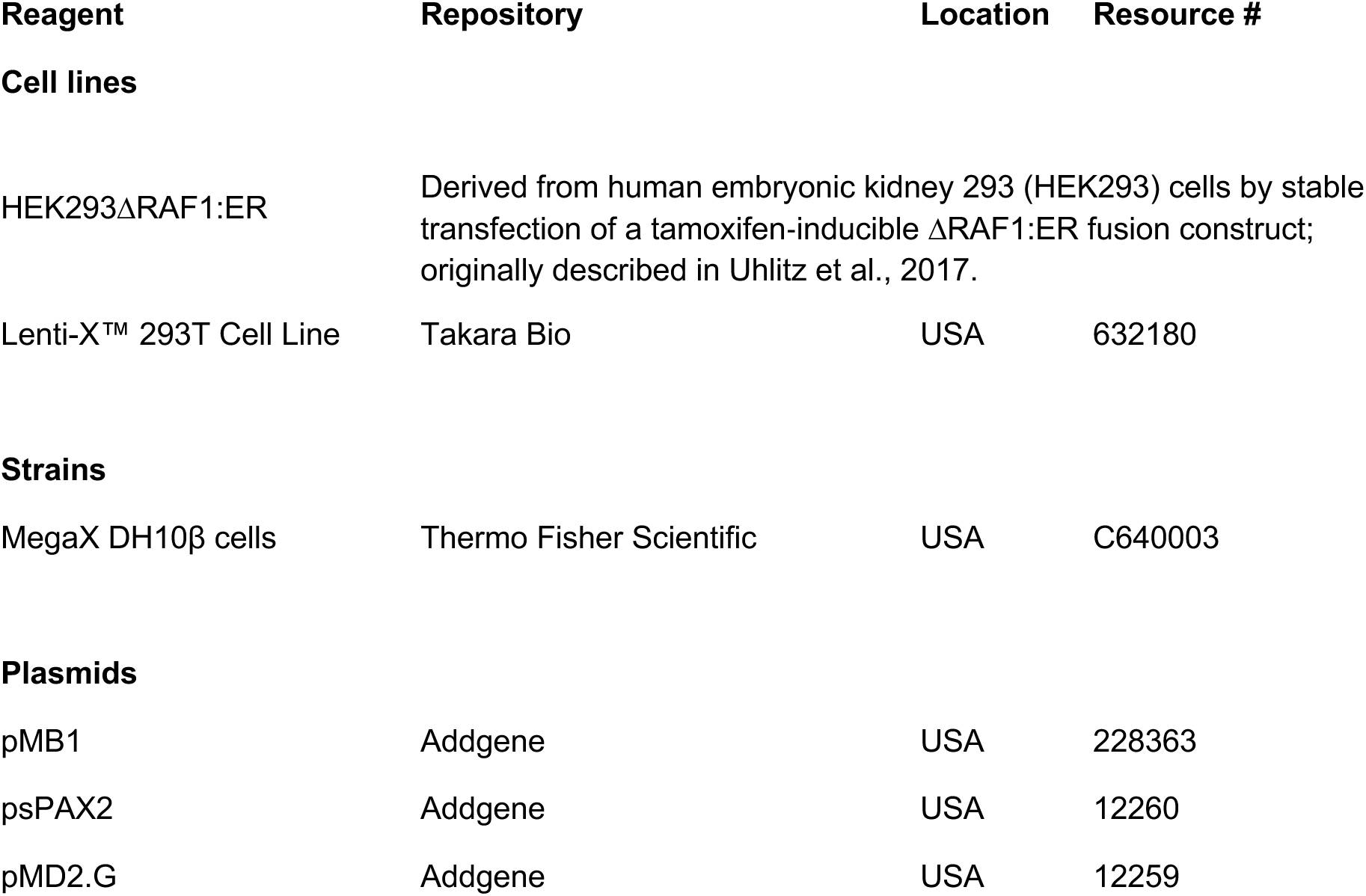

| Reagent | Repository | Location | Resource # |
| --- | --- | --- | --- |
| Cell lines |  |  |  |
| HEK293ΔRAF1:ER | Derived from human embryonic kidney 293 (HEK293) cells by stable transfection of a tamoxifen-inducible ΔRAF1:ER fusion construct; originally described in Uhlitz et al., 2017. |  |  |
| Lenti-X™ 293T Cell Line | Takara Bio | USA | 632180 |
| Strains |  |  |  |
| MegaX DH10β cells | Thermo Fisher Scientific | USA | C640003 |
| Plasmids |  |  |  |
| pMB1 | Addgene | USA | 228363 |
| psPAX2 | Addgene | USA | 12260 |
| pMD2.G | Addgene | USA | 12259 |

### Statistical analysis

For all experiments, the number of technical and/or biological replicates is listed in the figure legends or text. Pearson correlation was used to determine the *r* values. Wald-P values were adjusted using Bonferroni correction method. Statistical analyses were performed using GraphPad Prism 9 (GraphPad Software) or the R language programming environment.

### Vector Maps

The all-in-one Cas9 vector pMB1 (17), containing the 10x Genomics capture sequence 1, 5′-GCTTTAAGGCCGGTCCTAGCAA-3′, in the stem-loop of the Cas9-tracr sequence (18), referred to here as pMB1-10x, was used for the Perturb-seq experiments. The plasmid map is provided in GenBank format (Supplementary Data 1).

### HEK293ΔRAF1:ER cell culture

HEK293ΔRAF1:ER cells (19) containing a tamoxifen inducible fusion of the kinase domain of RAF1 (20, 21) reviewed in (20, 21) were cultured in complete DMEM low glucose without phenol red supplemented with 10% fetal bovine serum (Pan Biotech) and 1% antibiotics (pen/strep). Lenti-X 293T cells (Takara) were cultured in complete DMEM supplemented with 10% fetal bovine serum and 1% antibiotics (pen/strep).

### Bulk RNA-sequencing data generation and preprocessing

Total RNA was extracted with TRIzol. Sequencing libraries were prepared using Illumina TruSeq mRNA Library Prep Kit v2 and sequenced on Illumina HiSeq 2000. Raw reads were processed using the snakemake-workflows rna-seq-star-deseq2 pipeline, v1.2.0, and counts were subsequently analyzed using DESeq2 in R.

### Cas9 library design

Target genes were selected based on RNAseq data after induction of HEK293ΔRAF1:ER cells with 0.5 µM 4-hydroxytamoxifen (4OHT) at different time points (Fig. 1). The sgRNA library consisted of 4 sgRNAs per gene with 10 non-target control sgRNA and 10 safe cutters sgRNAs. The sgRNA sequences were selected from the Brunello genome wide library (22). 4 positive-control sgRNAs against the Raf-transgene were designed using CRISPick (22, 23) (Supplementary Table 1).

**Figure 1:**
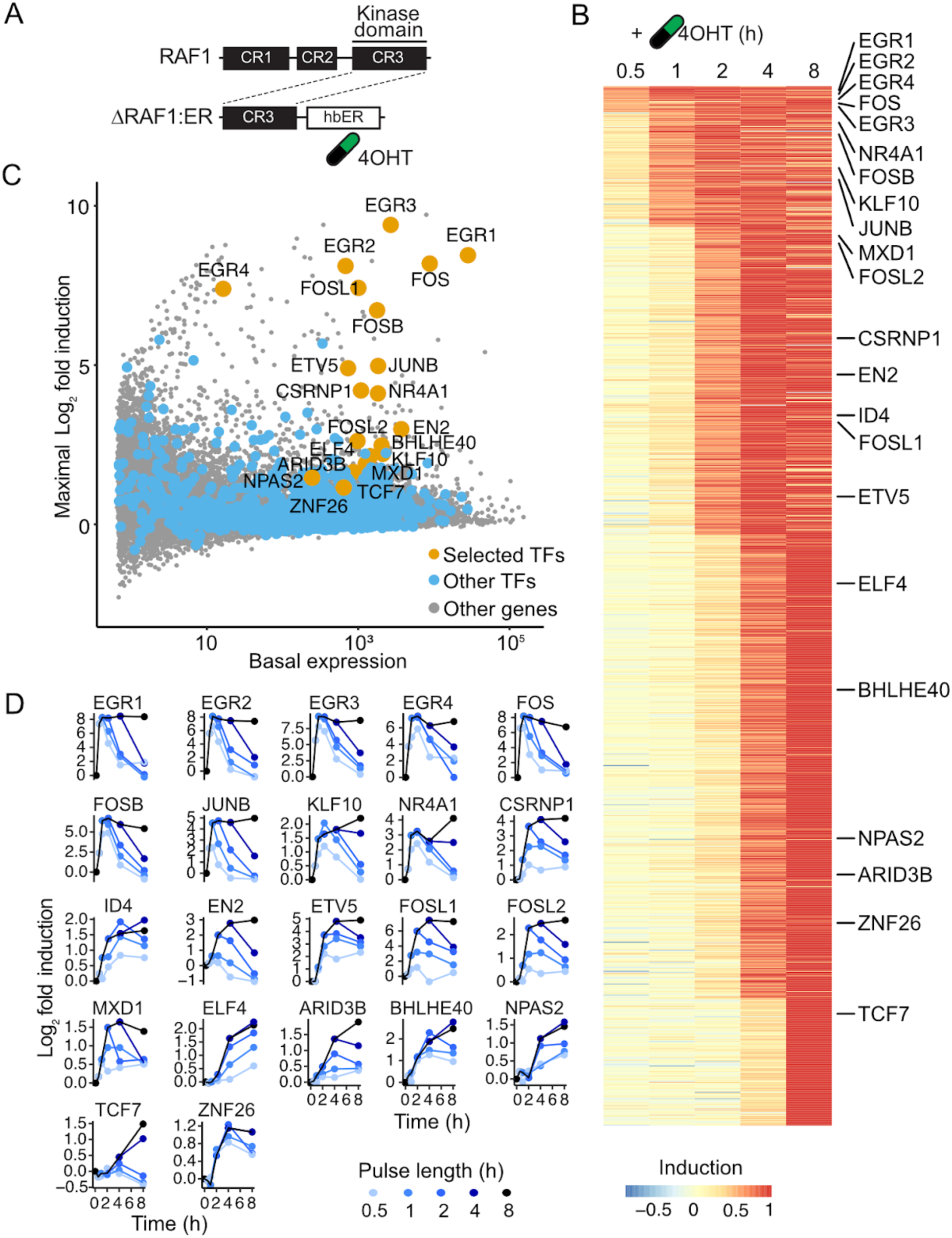
Transcriptional profiling of HEK293ΔRAF1:ER cells following RAF1 induction. **(A)** Schematic structure of the 4-hydroxytamoxifen-(4OHT)-inducible RAF1-CR3 kinase domain. **(B)** Maximum normalized log_2_ fold changes from bulk RNAseq analysis of significantly up-regulated genes (FDR<1%) after 0.5h, 1h, 2h, 4h and 8h. The 22 selected candidate TFs are indicated by their official gene symbols. **(C)** The maximal log_2_ fold expression change of the selected candidate TFs is plotted against their expression levels in non-induced HEK293ΔRAF1:ER cells. Orange: Selected TFs, Blue: Other TFs, Grey: Other genes. **(D)** The time-resolved log_2_ fold expression changes of candidate TFs are shown after the indicated pulse lengths of 4OHT-mediated RAF1 induction.

### Cas9 library cloning

The selected 20-nt target specific sgRNA sequences were cloned into the pMB1-10x library vector (Supplementary Data 1) by Gibson Assembly (24). sgRNA template sequences of the format: 5′-GGAGAACCACCTTGTTGG-(N)20-GTTTAAGAGCTAAGCTGGAAAC-3′ were synthesized in a pooled format on microarray surfaces (GenScript Biotech, Inc.). Oligo pools were PCR-amplified using Phusion Flash High-Fidelity PCR Master Mix (ThermoFisher Scientific) according to the manufacturers protocol with 1 ng/μL sgRNA template DNA, 1 μM forward primer (5′-GGAGAACCACCTTGTTGG-3′), and 1 μM reverse primer (5′-GTTTCCAGCTTAGCTCTTAAAC-3′) in 50 µL total volume. The following cycle numbers were used: 1× (98 °C for 3 min), 16× (98 °C for 1 s, 54 °C for 15 s, 72 °C for 20 s) and 1× (72 °C for 5 min). PCR products were purified using NucleoSpin columns (Macherey-Nagel). The library vector pMB1-10x was prepared by restriction digestion with AarI (Thermo Fisher) at 37 °C overnight. The digestion reaction was run on a 1% agarose gel followed by excision of the digested band and purification via NucleoSpin columns (Macherey-Nagel). 100 ng digested pMB1-10x and 2.4 ng amplified sgRNA library insert were assembled using Gibson Assembly Master Mix (NEB) in a 20 μL reaction for 30 min. The reaction was purified using P-30 buffer exchange columns (BioRad) that were equilibrated 5x with H_2_O and the eluted volume was transformed into 20 µL of MegaX DH10β cells (Thermo Fisher) by electroporation. *Escherichia coli* were recovered and cultured overnight in 100 mL LB medium with 100 μg/mL ampicillin. The plasmid library was extracted using Midiprep (Qiagen). In parallel, a fraction of the transformation reaction was plated and used to determine the total number of transformed clones. The coverage was determined to be 1,643x clones per sgRNA ensuring even representation of all library sgRNA sequences and their narrow distribution (Fig. 2B). The quality of the cloned sgRNA library was determined by NGS on an Illumina Miniseq (See below). MAGeCK (25) was used for library alignment. Narrow distribution of sgRNA sequences was confirmed with read counts for 96% of sgRNA sequences falling within a single order of magnitude.

**Figure 2:**
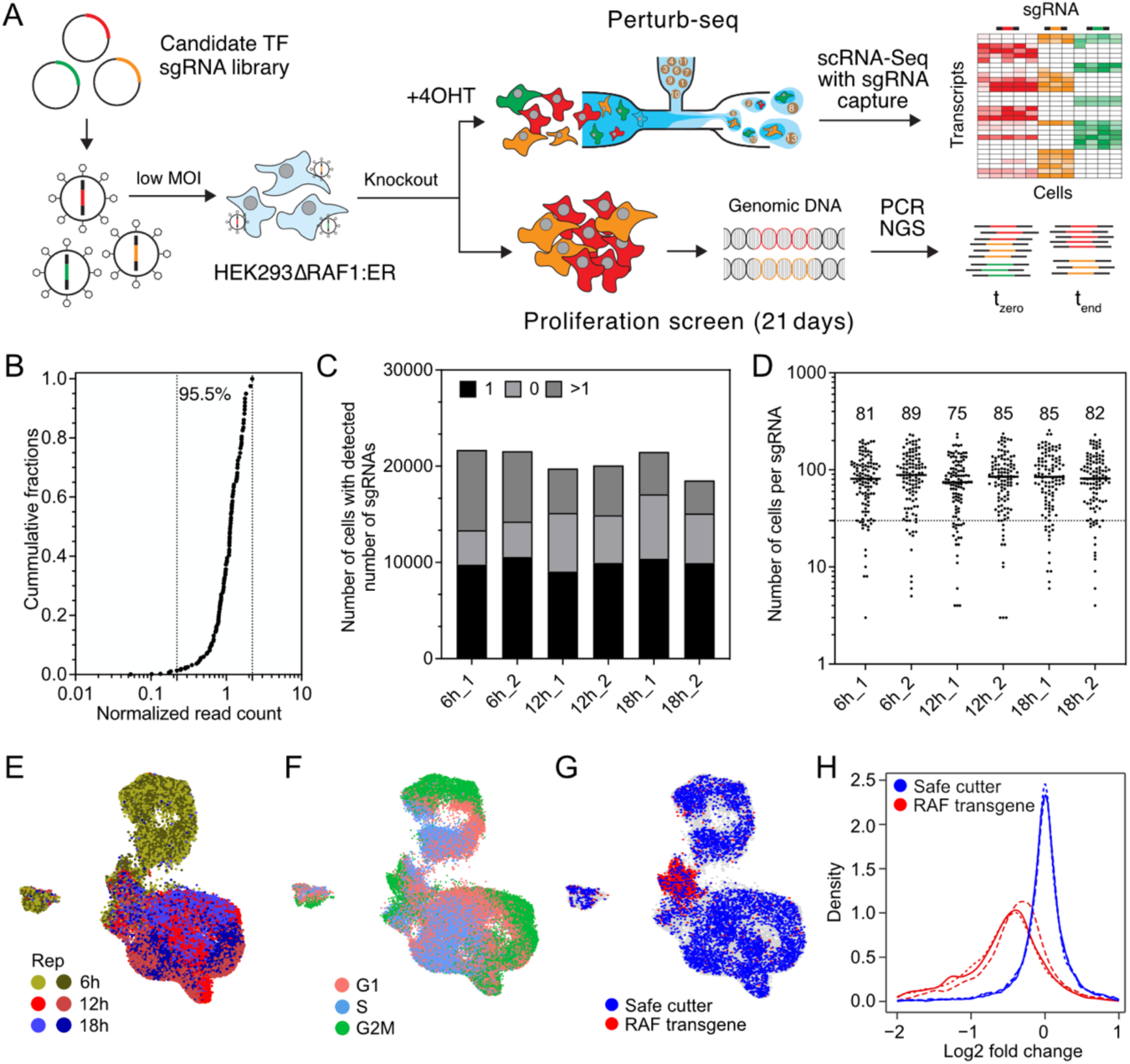
Perturb-seq and proliferation screens in HEK293ΔRAF1:ER. **(A)** Schematic of Perturb-seq screens (top panel) and proliferation CRISPR screens (bottom panel). **(B)** Distribution of the pooled sgRNA library used for all Perturb-seq and proliferation screens. **(C)** Total number of recovered cells and number of cells with 0, 1, and >1 sgRNAs detected in the respective Perturb-seq samples. **(D)** Median number and distribution of cells per sgRNA detected in the respective Perturb-seq samples. Dashed line at 30 cells per sgRNA indicates the cut-off cell number per sgRNA that was used for further analysis. **(E)** UMAP integration of Perturb-seq samples from three different time points with two replicates each. **(F)** Distribution of cell cycle phases G1, S, and G2M on the integrated UMAPs. **(G)** Distribution of the safe cutter sgRNA cells and the RAF1-knockout cells on the UMAP clusters. **(H)** Histograms of the log_2_ fold change of previously identified RAF/MAPK target genes (Fig. 1) between RAF1-knockout (red) and the safe cutter control cells (blue), relative to the non-target control cells. Dashed line: 6h time point, Dotted line: 12h time point, Solid Line: 18h time point. Perturb-seq screens were performed in duplicate.

### Lentivirus production

Lenti-X 293T cells (Takara) were seeded at 65,000 cells per cm^2^ in 10 mL media (DMEM, 10% FBS, 1% pen/strep) in a 10 cm dish and incubated overnight at 37 °C, 5% CO_2_. On the next day, 5 μg sgRNA library plasmid, 2 μg psPAX2 (Addgene #12260), 2 μg pMD2.G (Addgene #12259), and 36 μL Turbofect (Thermo Fisher) were mixed into 1.8 mL serum-free DMEM (Gibco), vortexed briefly, incubated for 20 min at RT, and added to the cells. At 48 and 72h post-transfection, the supernatant was harvested, passed through 0.45 um filters (Millipore), and 20x concentrated using LentiX Concentrator (Takara) according to the manufacturer’s instructions. Aliquots were stored at -20 °C.

### Direct capture Perturb-seq CRISPR screens

HEK293ΔRAF1:ER cells were transduced with lentivirally packaged sgRNA library at a multiplicity of infection (MOI) = 0.2 and 1,000x coverage in 2 replicates. The low MOI and high coverage were used to reduce the frequency of multiple-infected cells thus only one gene was knocked out in each cell and ensure even distribution of the sgRNA library. Cells were then cultured in DMEM low glucose without phenol red with 10% FBS (Pan Biotech) and 1% pen/strep (Sigma-Aldrich) in a 37 °C incubator with 5% CO_2_. 48h after transduction, transduced cells were selected with puromycin (2 μg/mL) for 96h. After selection, the top 20% mcherry positive cells were sorted 8 days post infection using a BD FACSAria II flow cytometer to increase sgRNA capture efficiency by the 10x Genomics Gel Beads. A total of 3 million cells were sorted. For the Perturb-Seq screen, 500,000 of the sorted cells were reseeded in full medium in 3 wells of a 12 well plate and incubated at 37 °C, 5% CO_2_. At day 10 post infection, the sorted cells were stimulated with 0.5 µM 4OHT (Sigma-Aldrich, H7904) for 6, 12, and 18h. After the incubation time, the cells were harvested followed by scRNA-Seq following the 10x Genomics Chromium Next GEM Single Cell 3’ Reagent Kits v3.1 (Dual Index) with Feature Barcode technology for CRISPR Screening protocol.

### Pooled proliferation CRISPR screen

The remaining 2.5 million sorted HEK293ΔRAF1:ER cells from day 8 post infection were reseeded in full medium in a 6 well plate and incubated at 37 °C, 5% CO_2_. On day 18 post infection, 0.5 µM 4OHT was added for 48h to induce the RAF1 transgene expression and apoptosis of the induced cells. After 48h, the dead cells were detached and removed with the medium. The living cells were reseeded in tamoxifen containing medium and incubated at 37 °C, 5% CO_2_ for 48h more to increase the cell selection efficiency. Aliquots of 500,000 cells from the 48h induction time point were taken. The cells were centrifuged, and the cell pellets were frozen down for later analysis via NGS.

### Genomic DNA extraction and PCR recovery of gRNA sequences

The genomic DNA (gDNA) was extracted from the HEK293ΔRAF1:ER cells using the Qiagen Genomic DNA extraction kit according to the manufacturer’s instructions. Two nested PCR reactions were performed to amplify the sgRNA cassette from the extracted gDNA. For the first PCR reactions, 5 μg gDNA, 0.3 μM forward (5′-GGCTTGGATTTCTATAACTTCGTATAGCA-3) and reverse (5′-CGGGGACTGTGGGCGATGTG-3′) primer, 200 μM dNTP mix, 1x Titanium Taq buffer and 2 μL Titanium Taq polymerase (Takara) were mixed in 50 µL total volume. The PCR reaction was run using the following cycles: 1x (94 °C, 3 min), 20x (94 °C, 30 s, 65 °C, 10 s, 72 °C, 20 s), 1x (68 °C, 2 min). For the second PCR reactions, 5 μL first-round PCR, 0.5 μM forward (5′-AATGATACGGCGACCACCGAGATCTACACACACTCTTTCCCTACACGACGCTCTTCCGATCTTCCCTTGGAGAACCACCTTGTTGG-3′) and reverse (5′-CAAGCAGAAGACGGCATACGAGAT-(N)_6_-GTGACTGGAGTTCAGACGTGTGCTCTTCCGATC-3′) primer where (N)_6_ is a 6 nt index for sequencing on the Illumina Miniseq platform, 200 μM dNTP mix, 1x Titanium Taq buffer and 2 μL Titanium Taq (Takara). PCR cycles were: 1x (94 °C, 3 min), 20x (94 °C, 30 s, 55 °C, 10 s, 72 °C, 20 s), 1x (68 °C, 2 min). The PCR product (325 bp) was purified from a 1% agarose gel via NucleoSpin columns (Macherey-Nagel). NGS was performed on an Illumina Miniseq using a MiniSeq Mid Output Kit (300-cycles) using paired-end 150 strategy according to the manufacturer’s instructions.

### Proliferation screen data analysis

The proliferation screen data analysis was performed using MAGeCK (25). In short, sgRNA read count files were computed from the raw CRISPR fastq files using the count function. The MAGeCK MLE command was then used to calculate the MAGeCK Beta score, Wald-P values, and false discovery rates for enrichment and depletion of each guide at day 20 and day 22 after tamoxifen induction compared to the plasmid library. Wald-P values were adjusted using Bonferroni correction method in R (Supplementary Table 2).

### scRNA-Seq screen data analysis

Cell Ranger (10x Genomics) Version 6.1.1 was used for scRNA-Seq data processing (https://support.10xgenomics.com/single-cell-gene-expression/software/pipelines/latest/using/count). Sequencing reads coming from the gene expression library were mapped to the GRCh38-1.2.0 genome reference compiled by 10x Genomics for Cell Ranger. Guide RNA reads were mapped simultaneously to a sgRNA feature reference. The combination of standard and targeted RNA-Seq was processed by pooling the fastq files and subsequent Cell Ranger analysis, thereby avoiding duplicate counts for the same molecules as reads with the same UMIs are collapsed. Count matrices were then used as input into the Seurat R package (26) to perform downstream analyses. Differential expression was called based on pseudo bulks using the R-library glmGamPoi version 1.10.2 (27).

### Gene expression library amplification (TAP-seq)

For the generation of suitable primers, we used the BAM file of an untargeted 10x run (6h 4OHT) in the TAP-seq R package (https://github.com/argschwind/TAPseq) and workflow that is delineated in the package vignette (16). Deviations from the TAP-seq workflow were as follows: (i) We only used the inner primer generation procedure, i.e. at 150-300bp from inferred poly(A) sites. (ii) Originally primers for one major poly(A) site per gene were generated, instead we generated primers for all top poly(A) sites amounting to >70% of the total poly(A) score (i.e. coverage) per gene. (iii) Additional filter steps on poly(A) level, i.e., removal of minor poly(A) sites in adjacent poly(A) signals within 100bp, and on primer level, i.e., removal of redundant primers within 500bp of poly(A) signal and manual filter of badly designed primer e.g., in non-expressed regions. The final primer list is provided in the Supplementary Table 3. Targeted primers were ordered from IDT in desalted format with 5’-GTGACTGGAGTTCAGACGTGTGCTCTTCCGATCT-3’ as PCR handle at 5’ ends. 100 µL PCR was performed as follows: 100 ng amplified cDNA (Step 2.3 of 10x Genomics 3’scRNA), 2.5 µL 100 µM Pooled targeted Primer Mix, 4 µL 10 µM partial Read 1(sequence from 10x Genomics manual, ordered as normal (desalted) primer from IDT 5’-CTACACGACGCTCTTCCGATCT-3’, 50 µL KAPA HiFi HS RM (Roche, KK2601). The PCR was performed with the following cycle numbers: 1x (95 °C, 3 min), 10x (98 °C, 20 s, 67 °C, 60 s, 72 °C, 60 s), 1x (72 °C, 5 min). The PCR product was cleaned using SPRIselect beads (Beckmann Coulter) as a double sided size selection (as described in the Tips & Best Practices Section of a typical 10x Genomics procedure) with 0.6x as 1^st^ SPRI and 1.2x as 2^nd^ SPRIselect steps. For adding indices, maximally 10 ng of the first cleaned PCR product were then mixed with 20 µL dual Index TT Set A from 10x genomics, 50 µL KAPA HiFi HS RM (Roche, KK2601) in a 100 µL reaction. The PCR was performed with the following cycle numbers: 1x (95 °C, 3 min), 1x (98 °C, 45 s), 10x (98 °C, 20 s, 54 °C, 30 s, 72 °C, 20 s), 1x (72 °C, 1 min). As the amplicons are bigger now, we changed the SPRI concentration to 0.55x as 1^st^ SPRI and 1.2x as 2^nd^ SPRIselect steps. Library quality was assessed using the TAPE station.

### EGR1 and FOS knockout clonal line production

HEK293ΔRAF1:ER single knockout clones for FOS and EGR1 genes were generated using CRISPR-Cas9. Two sgRNAs per gene were designed to target 500 bp sequences surrounding the 5’ end using CRISPOR (28). The sgRNAs and their reverse complements were synthesized and cloned into the px459 vector (29). For the EGR1 gene knockout, the following oligo sequences were used: 5’-CACCGGGCCATGTACGTCACGACGG-3’ and 5’-CACCGGGACAACTACCCTAAGCTGG-3’ targeting the promoter and the exon regions respectively. For the FOS gene knockout, the following oligos sequence was used to target the promoter region; 5’-CACCGGATTAGGACACGCGCCAAGG-3’ and the exon region, 5’-CACCGGAGAGAGGCTATCCCCGGCCG-3’. The oligo for FOS exon contains an added G at the 5’ end of the gRNA to facilitate U6 promotor mediated transcription. Cells were transfected with these vectors following the Lipofectamine 2000 (ThermoFisher Scientific) and selected with puromycin (250 ng/mL) for 36h starting 24h after transfection. Following selection, we performed clonal dilution. Cells were seeded in 96 well plates and wells with individual clones were screened via PCR to identify successful 5’ end deletions. Positive hits were further validated at RNA and protein levels using qPCR and Western blot, to confirm the absence of FOS and EGR1 gene expression.

Starting from single knockout clones of EGR1 and FOS, double knockout clones were generated using Alt-R™ S.p. Cas9 Nuclease V3 from IDT used with guides designed with the manufacturer’s design tool (https://eu.idtdna.com/site/order/designtool/index/CRISPR_CUSTOM) (Supplementary Table 4). Transfection was performed following the Lipofectamine CRISPRMAX (ThermoFisher Scientific), using less RNA amounts depending on the number of sgRNAs used per transfection. Two days after transfection, clonal dilution was performed. Isolation of gDNA was done with Quick-DNA-96 Kits (Zymo), for PCRs KAPA HiFi HS RM (Roche) and different primer sets spanning regions out and/or inside expected deletions were used. PCR product size was analyzed with TAPE Station from Agilent. Clones with successful deletions based on the PCR result were then analyzed using Western blots with antibodies against FOS or EGR1 and pERK antibody (Cell Signaling Technology) to identify clones that lack expression of the respective TFs and still induce MAPK signaling when the RAF1-CR3 kinase domain is activated with 4OHT. KO clones were cultivated in low glucose DMEM without Phenol red (Sigma-Aldrich) supplemented with stable Glutamine (PAN-Biotech) and FBS (PAN-Biotech).

### Bulk RNA-Seq

For bulk RNA sequencing of single and combinatorial EGR1 and FOS knockout clonal lines cells were treated with 0.5 µM 4OHT for 6h, 12h, and 18h. To account for differences in cell density, two solvent control wells were collected each at first and last lysing time of the treatments. RNA isolation was done with RNeasy Kits from Qiagen without any DNA elimination. RNA concentration was measured with the Implen nanophotometer and 250 to 450 ng per sample were used for Library preparation with QuantSeq 3’ mRNA-Seq V2 (Lexogen). For Index PCR, 18 cycles were used according to the manual. Upon quantification with Universal KAPA Library Quantification Kit for Illumina (Roche) Libraries were pooled and sent for sequencing.

For bulk RNA sequencing of TCF7 knockout cells after Wnt signaling activation, TCF7 and Safe-cutter knockout cells were made by transducing the cells with lentiviruses made from pMB1-sgTCF7-2 and pMB1-sgSafe-cutter-1 vectors without the 10x Genomics capture sequence 1 in the stem-loop of the Cas9-tracr sequence. The cells were selected with 2µg/mL puromycin and incubated for 10 days. After the incubation period, th ecells were treated with 10 µM CHIR99021 for 24 hours. The cells were then harvested and the total RNA was extracted using the RNeasy Mini Kit (Qiagen, 74106) according to the manufacturer’s instructions. The RNA was then quantified using a NanoDrop One and sent for bulk RNA-seq at Novogene UK.

### Quantitative real-time polymerase chain reaction (qRT-PCR)

HEK293ΔRAF1:ER EGR1, TCF7, and Safe-cutter knockout cells were generated as before. For the MAPK and Wnt signaling pathway activation the cells were treated with 0.5 µM of 4OHT for 12 h and 10 µM of CHIR99021 for 24 h. For the EGR1-TCF7 positive feedback loop the cells were then treated with 0.5 µM of 4OHT for 1 h while the cells for checking the TCF7-EGR1 positive feedback loop were treated with 0.5 µM of 4OHT for 12 h. Finally, for investigating the effect of MAPK and Wnt signaling activation on DKK1, the cells were treated with 0.5 µM of 4OHT for 12 h and 10 µM of CHIR99021 for 6 hour either separately or in combination.

Total RNA was isolated for qRT-PCR with a RNeasy Mini Kit (Qiagen, 74106) according to the manufacturer’s instructions, and quantified using a Nanodrop One. cDNA was prepared from total RNA using a RevertAid H Minus First Strand cDNA Synthesis Kit (ThermoFischer Scientific, K1632). Quantitative PCR was performed using ORA qPCR Green ROX L Mix (highQu, QPD0105) using a LightCycler® 480 System (Roche). Analysis was performed using the 2^−ΔΔCt^ method. GAPDH was used as a housekeeping gene. The following primers were used: GAPDH_for: CTGGTAAAGTGGATATTGTTGCCAT, GAPDH_rev: TGGAATCATATTGGAACATGTAAACC, EGR1_for: CTTCAACCCTCAGGC GGACA, EGR1_rev: GGAAAAGCGGCCAGTATAGGT, TCF7_for: CTGACCTCTCTGGCTTCTACTC, TCF7_rev: CAGAACCTAGCATCAAGGATGGG, AXIN2_for: CAAACTTTCGCCAACCGTGGTTG, AXIN2_rev: GGTGCAAAGACATAGCCAGAACC. DKK1_for: CACACCAAAGGACAAGAAGG, and DKK1_rev: CAAGACAGACCTTCTCCACA.

### Re-analysis of scRNA-seq of colon organoids

Reanalysis of previously published single cell RNA-Sequencing data of colon organoids (30) was performed as follows. The two raw count matrices were first demultiplexed using the HTODemux algorithm from Seurat and subsequently merged. Data was then subseted to cells with more then 2000 features and less then 20% mitochondrial read. Only data for the lines NCO (normal), A (APC), AK (APC and KRAS) under the condition N (Noggin, i.e. without WNT/R-spondin and EGF) were used. Using AddModuleScore, the TCF7 signature (all significant genes of the perturb-seq screen with p_adj<0.1 and log2-fold change < 0.1 for the TCF7 guides), the TCF7 signature derived from the screen was scored. Raw data is available at: https://doi.org/10.5281/zenodo.7817521 (30).

### Processing of bulk, multiplexed QuantSeq data

bcl2fastq (v2.20.0 by Illumina) was used to demultiplex and convert raw sequencing data to fastq files. We designed a Snakemake (v7.18.2) workflow in which BBMap’s BBDuk (v39.01) was used to trim adapters, STAR (v2.7.10b) to align reads to the GENCODE GRCh38.p13 (v39) geneset, umitools (v1.1.4) to extract and deduplicate UMIs, and subread’s featureCounts (v2.0.6) to count mapped reads on gene level.

## RESULTS

### Identification of transcription factors up-regulated by RAF1-induction

To identify transcripts that are up-regulated by RAF-MAPK activation, we used a previously established HEK293 cell line, termed HEK293ΔRAF1:ER, in which a tamoxifen-inducible RAF1-CR3 kinase domain was introduced (20). Consequently, RAF1 activity can be precisely regulated, allowing the identification of RAF1-MAPK response genes (Fig. 1A). In contrast to cell culture systems stimulated by growth factors, this model of RAF-MAPK signaling is activated independently of the upstream G-protein RAS, thus minimizing pathway divergence and feedback mechanisms. We induced RAF activity with 4-hydroxytamoxifen (4OHT) over periods of 0.5 to 8h and monitored the changes in the transcriptome of the cells via bulk RNA-sequencing (RNA-Seq). Over the full time series, we detected a total of 1,142 significantly up-regulated genes (Fig.1B). From this dataset, 22 transcription factors (TFs) that were up-regulated at different time points after RAF1 induction were selected for further analysis. The basal expression levels of identified candidate TFs varied over several orders of magnitude and their level of up-regulation upon RAF1 induction was independent of their basal expression level (Fig.1C).

Pulsed induction of RAF1 (Fig. 1D) further allowed the categorization of the candidate TFs into three distinct response classes. The first class comprises classical immediate early genes that are rapidly induced upon signal induction and whose mRNAs are rather short lived, resulting in rapid decay after the pulses ended. Examples of these transcripts are the rapidly induced EGR gene family members, FOS, FOSB, and JUNB, which increased to maximum mRNA levels directly after induction and decreased quickly after 4OHT removal. The second class comprises rather rapidly induced, long-lived transcripts that are induced within the first 1-2h and remain high even after the pulse has ended, termed immediate late genes. These include, for instance, the transcripts of FOSL1 and FOSL2. A third class of transcription factors is only induced with delay. For several of those transcripts, the response is limited to long pulses of induction. For instance, TCF7 is only induced after 4h and reaches a maximum response after 8h of induction. We furthermore calculated the time when the transcripts reach half-maximal expression by interpolating between the measured time points. We found that the selected transcripts cover half-maximal induction times between 20 min and 5h (Table 1).

**Table 1:** Selected candidate genes and their classification based on their response time to RAF1 induction. (**8**). IEG = immediate-early genes, ILG = immediate-late genes, DEG = delayed-early genes. SRG = secondary response gene.

| Gene Symbol | Name | Class (8) | Half-maximal induction (h:min) |
| --- | --- | --- | --- |
| EGR1 | Early Growth Response 1 | IEG | 0:17 |
| FOS | Fos Proto-Oncogene, AP-1 Transcription Factor Subunit | IEG | 0:19 |
| EGR2 | Early Growth Response 2 | IEG | 0:22 |
| EGR3 | Early Growth Response 3 | IEG | 0:25 |
| EGR4 | Early Growth Response 4 | ILG | 0:29 |
| JUNB | JunB Proto-Oncogene | IEG | 0:33 |
| FOSB | FosB Proto-Oncogene | IEG | 0:37 |
| NR4A1 | Nuclear Receptor Subfamily 4 Group A Member 1 | ILG | 0:47 |
| KLF10 | KLF Transcription Factor 10 | DEG | 0:49 |
| ID4 | Inhibitor Of DNA Binding 4 | DEG | 1:05 |
| MXD1 | MAX Dimerization Protein 1 | DEG | 1:13 |
| FOSL1 | FOS Like 1 | ILG | 1:17 |
| CSRNP1 | Cysteine And Serine Rich Nuclear Protein 1 | SRG | 1:19 |
| FOSL2 | FOS Like 2 | ILG | 1:23 |
| ETV5 | ETS Variant Transcription Factor 5 | IEG | 1:31 |
| EN2 | Engrailed Homeobox 2 | SRG | 1:37 |
| ZNF26 | Zinc Finger Protein 26 | DEG | 2:09 |
| BHLHE40 | Basic Helix-Loop-Helix Family Member E40 | DEG | 2:15 |
| ARID3B | AT-Rich Interaction Domain 3B | SRG | 3:03 |
| ELF4 | E74 Like ETS Transcription Factor 4 | SRG | 3:10 |
| NPAS2 | Neuronal PAS Domain Protein 2 | SRG | 3:16 |
| TCF7 | Transcription Factor 7 | SRG | 5:08 |

The selected transcription factors exhibit distinct kinetics in the RAF1-induced transcriptional response, encompassing transcripts previously classified as immediate-early (IEG), immediate-late (ILG), delayed-early (DEG), and secondary response genes (SRG) (8). While these induction dynamics suggest whether a factor acts early or late in the RAF response, they do not reveal whether, or how, the induced TFs functionally contribute to transcriptional network regulation downstream of RAF–MAPK. To systematically define the transcripts controlled by the 22 TFs listed in Table 1, we employed Perturb-seq (14, 15).

### Perturb-seq reveals the transcriptional targets of RAF1-induced TFs

To identify the transcriptional targets of the selected 22 candidate TFs, we performed pooled CRISPR/Cas9 screens with single cell RNA-Seq read-out, via direct-capture Perturb-seq (18), as well as pooled CRISPR screens with proliferation read-out (Fig. 2A). For that purpose, we designed and cloned a pooled sgRNA library, targeting the 22 candidate TFs with 4 sgRNAs each, 10 non-target control and 10 safe cutter sgRNAs that cut in gene-free regions of the genome to control for potential DNA double-strand break induced side-effects. In addition, the library contained 4 sgRNAs against the 4OHT-inducible RAF1 transgene as a positive control. We confirmed narrow library distribution with more than 95% of the sgRNAs read counts falling within a single order of magnitude (Fig. 2B). The sgRNA library was then transduced into HEK293ΔRAF1:ER cells at low multiplicity of infection (MOI = 0.2) to ensure the introduction of maximum one sgRNA expression cassette per cell. Following a 10-day period to allow for CRISPR/Cas9 gene editing (Fig. S1), RAF1 was induced by 4OHT for 6h, 12h, and 18h, respectively, to capture time-resolved transcription profile changes from each of the 22 perturbed candidate TFs. Cells were then analyzed via single-cell RNA sequencing (scRNA-seq) using “Feature Barcode technology” (10X Genomics).

As expected, the low MOI used for transduction resulted in the detection of one sgRNA in the majority of captured cells (Fig.2C). The detection of more than one sgRNA per cell is most likely explained by multiple cells being captured in the same emulsion droplet (Fig.2C). Cells with >1 or 0 detectable sgRNA were omitted from further analysis, resulting in approximately 10,000 analyzable cells per sample expressing exactly one sgRNA and a median sgRNA UMI count of 26-84 (Fig. S2A). The median number of analyzable cells per sgRNA, varied between experiments, ranging from 75 to 89 cells (Fig. 2D) and sgRNAs with less than 30 analyzable cells were excluded from further analysis. Taken together, these results demonstrate the high technical quality of the generated Perturb-seq data, which formed the basis for further analyses.

### Quality control of Perturb-seq screen data

Integration of UMAPs from Perturb-seq experiments at 6, 12, and 18 hours after RAF1 induction revealed that cells from the 12- and 18-hour time points clustered separately from those at 6 hours (Fig. 2E). This indicates that major transcriptional changes occur between 6 and 12 hours, highlighting the temporal dynamics of the RAF1 response. Importantly, cells within each UMAP cluster were uniformly distributed across G1, S, and G2/M phases, ruling out cell cycle heterogeneity as a source of variation (Fig. 2F). To validate the system, we compared RAF1-transgene knockout cells to safe-cutter control cells. RAF1 knockouts formed a distinct cluster, clearly separated from the controls (Fig. 2G), confirming the absence of a transcriptional response to RAF1 induction in these cells. Consistently, pseudo-bulk analysis of the previously identified RAF–MAPK targets (Fig 1B) showed a normal distribution centered around zero in safe-cutter controls, whereas RAF1 knockouts displayed a clear shift toward negative log2 fold changes (Fig. 2H). These findings demonstrate that the transcriptional changes observed in control cells are specifically driven by RAF1 activation and not by Cas9-induced double-strand breaks. Examination of individual RAF1 sgRNAs confirmed that three guides were effective, with 12–37% of RAF1-induced genes (identified by bulk RNA-seq) significantly deregulated (Fig. S2B). The fourth sgRNA was represented in fewer than 30 cells and was excluded from analysis. Together, these results verify that the Perturb-seq screens were robust, with positive and negative controls performing as expected.

### Proliferation CRISPR screens after candidate TF perturbation

We next asked whether knockout of individual transcription factors affects cell growth, using a CRISPR/Cas9 proliferation screen performed in HEK293ΔRAF1:ER cells with and without RAF induction (Fig. 2A, S3). Most TF knockouts showed no significant depletion or enrichment compared to non-targeting controls. Guides targeting FOS and EGR2 were significantly depleted under both conditions, consistent with their known roles in cell fitness (31, 32). In addition, knockouts of ELF4, FOSL1, FOSB, and EGR3 also showed significant depletion, though to a lesser extent. By contrast, knockout of the ΔRAF1:ER transgene was strongly enriched, but only upon RAF activation, in line with previous reports that prolonged RAF–MAPK activation in HEK293ΔRAF1:ER cells induces apoptosis (19). Overall, this screen demonstrates that TF knockouts do not cause major growth disadvantages, ensuring sufficient guide representation for subsequent Perturb-seq experiments (Fig. S3).

### A modified TAP-seq approach enhances the sensitivity of Perturb-seq screens

Perturb-seq screens are powerful but limited by sequencing costs, low sensitivity for weakly expressed genes, and small effect sizes. To address this, we adapted Targeted Perturb-seq (TAP-seq; (16)) by using PCR amplification directly from pre-amplified cDNA libraries, increasing sensitivity without loss of specificity (Fig. S4A). This modification reduces contamination risk, allows retrospective amplification of new gene panels, and was applied here to 140 RAF1-responsive genes (Fig. S4B). The approach maintained specificity in safe-cutter controls while substantially increasing the proportion of significantly deregulated genes in RAF1 knockouts and candidate TF perturbations, including EGR1 and FOS (Fig. S4C).

### Number of deregulated target genes varies greatly between TFs

After having established a modified TAP-seq protocol, we combined the targeted and untargeted Perturb-seq data to optimally analyze the 140 selected transcripts at high sequencing depth. In addition, it covers the entire transcriptome from the standard gene expression library. Figure 3 summarizes the results of a pseudo-bulk analysis of the combined targeted and untargeted Perturb-seq screens at 6h, 12h, and 18h after RAF1 induction, analyzed separately for each sgRNA.

**Figure 3:**
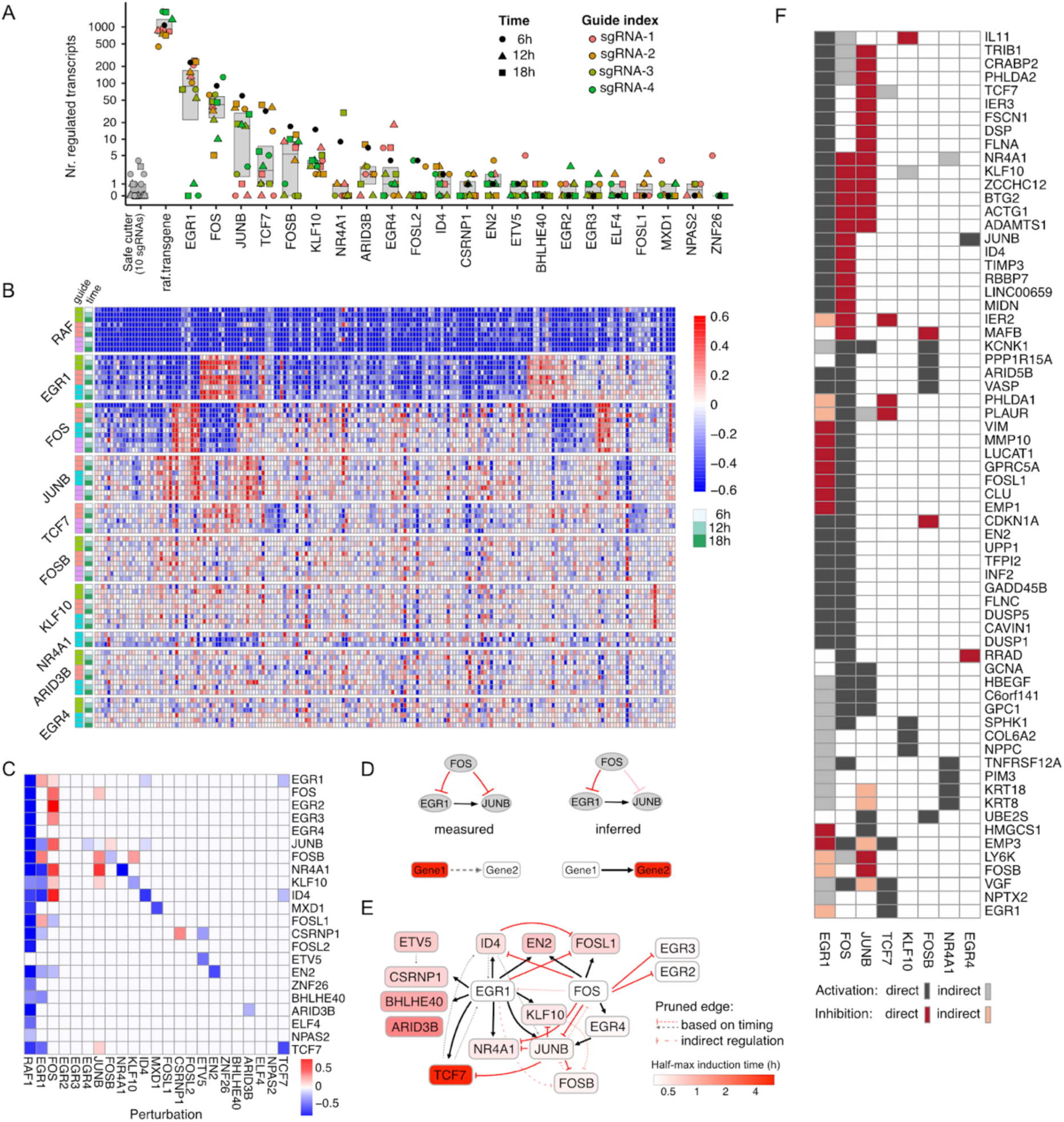
Summary of Perturb-seq screen results and model of transcriptional interactions between candidate TFs. **(A)** Number of significantly deregulated transcripts (adjusted p-value <0.05) following perturbation of the indicated candidate TF separated by sgRNAs and time points. **(B)** Heatmap of the log_2_ fold change of the 140 genes included in the modified TAP-seq for the perturbations with the strongest transcriptional response. The results from perturbed target genes separated by sgRNAs and time points are shown. Negative values indicate lower and positive values higher target transcript levels in the perturbed cells relative to cells expressing non-target control sgRNAs. **(C)** Core heatmap of candidate TF expression changes following the perturbation of all 22 TFs and RAF1, showing the log_2_ fold change upon perturbation for the significantly differentially expressed TFs (adjusted p-value <0.05). **(D)** Example for the removal of edges from coherent feed forward loops: The measured perturbation data in A is compatible with a feed-forward loop, where FOS inhibits JUNB directly and indirectly, but it is also compatible with a cascade, where FOS inhibits JUNB via EGR1 only. To generate the most parsimonious network, we removed the feed forward loop from FOS to JUNB in the inferred network and termed it “indirect”. Edges from late (red) to early (white) induced genes are dotted lines while the opposite orientation is represented by solid lines. **(E)** De-novo model of the TF core network showing directional interactions between all perturbed TFs with inferred interaction type: Black = activating, Red = inhibiting. Transparent edges indicate the removed feed forward loops. Transcription factors are color-coded by their half-maximal induction time. **(F)** Summary of target genes that are directly or indirectly co-regulated by more than one candidate TF.

Consistent with Fig. S4C, negative control sgRNAs produced no significant deregulation, whereas RAF1 knockouts affected 500–1,000 genes across all time points (Fig. 3A). The effects of TF perturbations varied widely: EGR1 knockouts deregulated hundreds of targets, while more than half of the investigated TFs had little or no effect. Heatmap analysis (Fig. 3B) confirmed that RAF1 loss blocked induction of its targets, with most changes evident by 6–12h and minimal additional effects at 18h, consistent with known RAF1 induction kinetics (1).

### Perturb-seq results are highly reproducible

To technically validate the Perturb-seq findings, we analyzed EGR1 and FOS single and double knockout clones in HEK293ΔRAF1:ER cells by bulk RNA-seq. The results closely resembled the Perturb-seq screen results, showing consistent directionality of regulation and strong correlations for the 140 TAP-seq transcripts (r = 0.69–0.8; Fig. S5A,B). Combinatorial EGR1-FOS knockouts clustered with the single knockout of FOS across both methods, suggesting that FOS is epistatic to EGR1 in the regulation of the RAF-MAPK response (Fig. S5C). Although the global profile of the double knockout resembled FOS loss, it also uncovered a synergistic role of EGR1 and FOS in upregulating EGR2, EGR3, and EGR4, which were strongly induced only in the double knockout (Fig. S5D).

### De-novo construction of a TF core network identifies EGR1 and FOS as dominant regulators of the RAF-MAPK transcriptional response

Because target gene induction was similar across time points, we combined sgRNAs and time points to define overall fold changes and applied Fisher’s method to identify significantly deregulated targets, resulting in a core heatmap of candidate TFs (Fig. 3C). Most TF knockouts also reduced their own transcript levels, consistent with nonsense-mediated decay of Cas9-induced stop-gain mutations (33). In contrast, EGR1, FOS, and CSRNP1 were upregulated, likely reflecting auto-regulatory feedback in which loss of protein function enhances transcription of mutant mRNA. Within the core network, EGR1 emerged as a central activator and FOS as a central inhibitor, often acting in opposition on shared targets such as JUNB, KLF10, ID4, and NR4A1. FOSL1 was oppositely co-regulated, while EN2 was co-activated by both TFs. These findings highlight EGR1 and FOS as orthogonal regulators, orchestrating the RAF1-driven transcriptional response.

To assess biological plausibility, we integrated time-series RNA-seq data (Fig. 1), removing nine edges where target activation preceded upstream TF induction, most likely representing indirect inhibitory effects. The final network also included a putative feed-forward loop (EGR1 → JUNB → FOSB), which could not be distinguished from a simpler model based on perturbation data alone. Applying parsimony, we excluded such ambiguous loops, yielding a refined core network (Fig. 3E). We then extended the analysis to target genes beyond the 22 candidate TFs. Figure 3F presents target genes co-regulated by the candidate TFs with the broadest regulatory impact, including EGR1, FOS, JUNB, TCF7, KLF10, FOSB, NR4A1, and EGR4.

### EGR1 interacts in a positive feedback loop with TCF7

Among all nodes in the core network (Fig. 3E), only one positive feedback loop was identified - between EGR1 and TCF7. EGR1 is a well-established downstream effector of MAPK signaling induced by 4OHT in HEK293ΔRAF1:ER cells, whereas TCF7, also induced by 4OHT (Fig. 1), is best known as a Wnt pathway regulator. This suggested that TCF7 might function as a nexus linking MAPK- and Wnt-driven transcriptional programs through its interaction with EGR1. To test this, we activated Wnt signaling in HEK293 cells with the GSK3β inhibitor CHIR99021, which stabilizes β-catenin and activates canonical Wnt signaling (Fig. 4A). Expression analysis confirmed pathway specificity: 4OHT strongly induced EGR1 (188-fold) and TCF7 (5.8-fold) but only marginally affected the Wnt marker AXIN2 (1.3-fold), indicating no activation of Wnt signaling (Fig. 4B). In contrast, CHIR99021 robustly upregulated AXIN2 (8-fold) and TCF7 (3-fold), consistent with Wnt activation, and also increased EGR1 expression (5.2-fold). We next validated the feedback loop by measuring EGR1 and TCF7 after their respective knockouts. TCF7 loss modestly but significantly reduced EGR1 expression, while EGR1 knockout lowered TCF7 levels (∼0.7-fold), confirming mutual activation (Fig. 4C).

**Figure 4:**
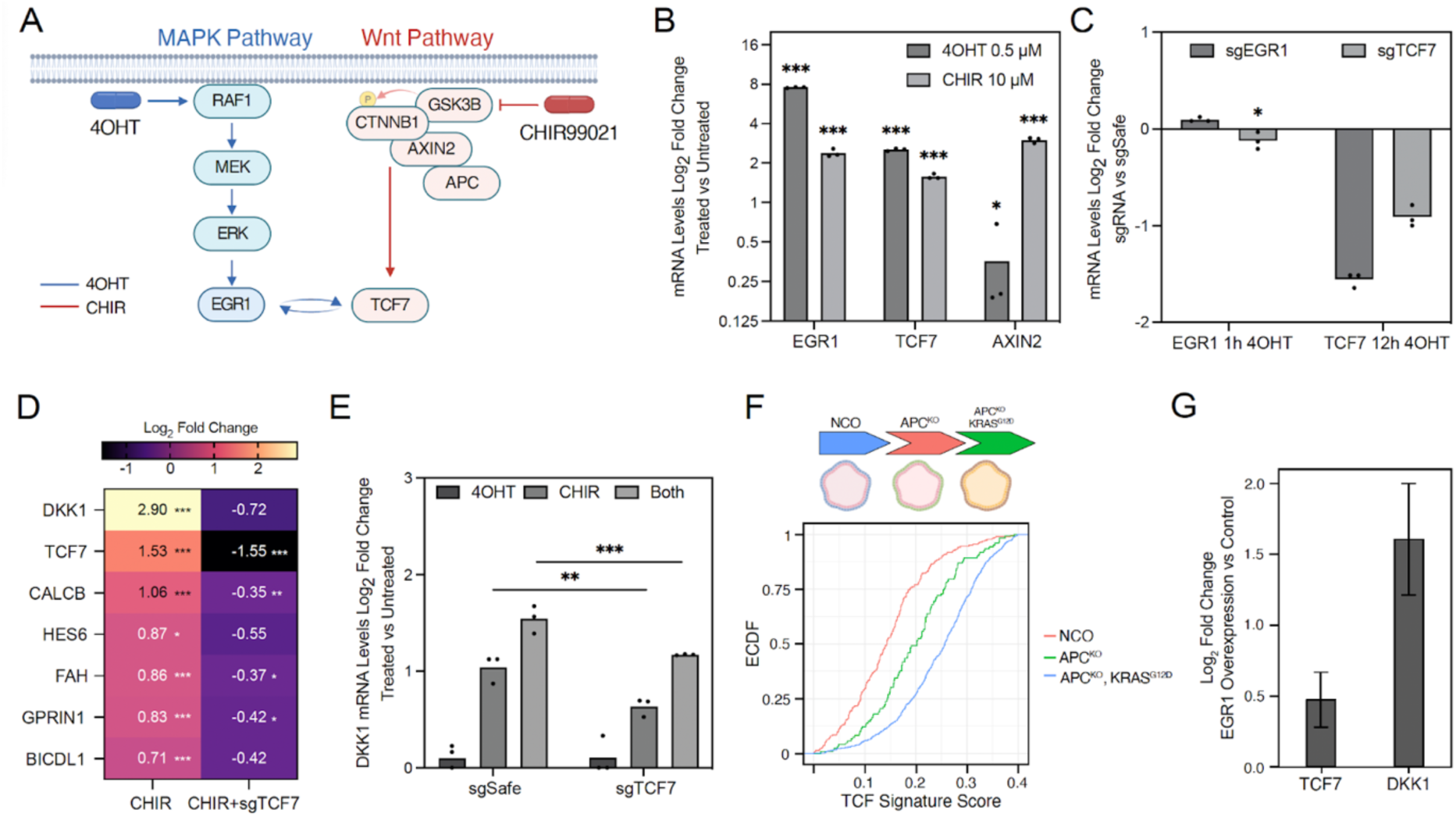
Characterization of MAPK and Wnt signaling pathways cross-talk through TCF7. **(A)** Simplified schematic of the MAPK- and Wnt-signalling pathways, including the EGR1-TCF7 positive feedback loop. RAF-MEK-ERK pathway is activated by 4OHT treatment. Wnt signaling is activated by CHIR99021 GSK3B inhibitor. Created in BioRender. Böttcher, M. (2026) https://BioRender.com/8k6fcao. **(B)** RT qPCR measurement of EGR1, TCF7, and AXIN2 mRNA levels after treatment of HEK293ΔRAF1:ER cells with 0.5 µM 4OHT or 10 µM of CHIR99021. Values represent the mean of biological replicates (n=3). **(C)** Validation of the EGR1-TCF7 positive feedback loop identified in the Perturb-seq screens through RT qPCR measurement of EGR1 and TCF7 mRNA level fold change after perturbation of EGR1 and TCF7. Values represent the mean of biological replicates (n=3). **(D)** Heatmap of transcriptional changes detected via bulk RNA-Seq from cells treated with 10 µM of CHIR99021 with and without TCF7 knockout. Log_2_ Fold Change values are calculated in reference to untreated cells. Values represent the mean of biological replicates (n=3). **(E)** RT qPCR measurement of DKK1 mRNA levels after treatment of the cells with 0.5 µM 4OHT for 12 h, 10 µM CHIR99021 for 6 h, or both. Values represent the mean of biological replicates (n=3). **(F)** TCF gene signature score showing the differences in expression of a set of differential expressed genes identified in the Perturb-seq screens after TCF7 knockout in colon organoids carrying inactivating APC and/or activating KRAS^G12D^ mutations. NCO = Normal Control cells with APC^wt^ and KRAS^wt^. **(G)** Reanalysis of a RNA-Sequencing dataset (GSE209899) showing DKK1 and TCF7 mRNA log_2_ fold changes after overexpression of EGR1 in Hela cells. Values represent the mean of biological replicates (n=3). * Adj. p-value <0.05, ** Adj. p-value <0.01, *** Adj. p-value <0.001.

### TCF7 serves as a nexus linking MAPK- and Wnt-driven gene expression

Bulk RNA-seq after CHIR99021 treatment revealed TCF7-dependent induction of canonical Wnt targets, including DKK1, CALCB, HES6, FAH, GPRIN1, and BICDL (Fig. 4D). RT-qPCR confirmed that DKK1 was induced by CHIR99021 (2-fold) and synergistically enhanced by combined MAPK and Wnt activation (2.9-fold) (Fig. 4E). This cooperative effect was reduced upon TCF7 knockout, confirming its role as a mediator of MAPK–Wnt cross-talk. To generalize these findings, we examined the expression of the identified TCF7 signature genes from the Perturb-seq data (Supplementary Table 5) in colon organoids and compared these to organoids with an APC knock-out and an additional KRAS^G12D^ mutation. TCF7 signature genes was upregulated by APC loss and further increased upon KRAS^G12D^ activation, confirming pathway synergy in a disease-relevant context (Fig. 4F). Finally, reanalysis of a public dataset (34) (GEO GSE209899) showed that EGR1 overexpression in HeLa cells not only induced TCF7 but also the bona fide Wnt target DKK1, further supporting the EGR1-TCF7 feedback loop as a mechanism of MAPK–Wnt integration (Fig. 4G).

## DISCUSSION

The MAPK signaling cascade has long served as a model for how extracellular cues are translated into defined transcriptional responses. Immediate-early and delayed response programs have been described in depth (2, 5, 7, 8), yet how early transcription factors interact to orchestrate downstream programs has remained poorly understood. Here, we applied targeted Perturb-seq to systematically dissect the RAF–MAPK transcriptional network, uncovering key features of its architecture.

To reconstruct the network topology, we applied a parsimony principle, removing coherent feed-forward loops to derive the simplest model consistent with the data, similar to ARACNE-based approaches (35). While coherent feed-forward loops are known to exist in transcriptional networks and may possess biologically significant functions (36), their removal aided in deriving a more streamlined network structure. Our analysis identified EGR1 and FOS as dominant hubs in the transcriptional RAF-MAPK response. Both factors control largely overlapping sets of target genes, often in an orthogonal manner. The observation that the EGR1–FOS double knockout largely resembled FOS loss suggests that FOS is epistatic to EGR1 for most targets, yet the synergistic regulation of EGR2-4 indicates a more nuanced interplay. This highlights how transcriptional networks combine dominant and redundant interactions to fine-tune cellular responses.

The most striking finding however, was the positive feedback loop between EGR1 and TCF7. EGR1 is a canonical MAPK effector (37), while TCF7 is classically linked to Wnt/β-catenin signaling (38). Their mutual activation creates a direct transcriptional bridge between MAPK and Wnt pathways. Functional assays confirmed that combined MAPK and Wnt activation synergistically enhanced Wnt target gene expression, including DKK1, a well-established Wnt feedback regulator (39). This transcriptional integration was not restricted to HEK293 cells but extended to colon organoids carrying APC and KRAS mutations, where dual pathway activation further boosted a TCF7 signature expression. Such synergy may help explain why MAPK and Wnt mutations frequently co-occur in colorectal and pancreatic cancer.

Our results indicate that the interplay between these two transcription factors provides a mechanism by which MAPK signaling can reinforce Wnt pathway activity, potentially enhancing oncogenic transcriptional programs. Kim et al. identified a hidden positive feedback loop caused by crosstalk between the Wnt and ERK pathways, suggesting that such feedback mechanisms can contribute to sustained activation of both pathways in cancer (40). The identification of the EGR1-TCF7 positive feedback loop provides a specific example of this phenomenon, where EGR1, a downstream transcription factor of the RAF-MEK-ERK MAPK pathway, and TCF7, a key component of the Wnt pathway, mutually reinforce each other’s expression.

Taken together, these results add a new layer to MAPK–Wnt cross-talk, which has largely been studied at the receptor, kinase, or β-catenin level (41). Here, we show that transcriptional network design encodes inter-pathway communication, with TCF7 acting as a nexus integrating MAPK and Wnt inputs. Such coupling may contribute to oncogenic robustness and therapy resistance, for example to MEK inhibition in KRAS-driven cancers (42). In conclusion, our work defines the architecture of the RAF-MAPK network, identifies EGR1 and FOS as central hubs, and establishes TCF7 as a key transcriptional link between MAPK and Wnt signaling.

## AUTHOR CONTRIBUTIONS

Ghanem El Kassem: Optimized and executed Perturb-seq and proliferation screens, cloned sgRNA library, performed RT qPCR experiments, manuscript writing. Anja Sieber: Designed and conducted targeted Perturb-seq experiments, transcriptome experiments, and generated and analyzed double KO lines. Bertram Klinger: Designed targeted Perturb-seq experiment. Florian Uhlitz: Performed time-series and pulsed transcriptome experiments. David Steinbrecht: Performed transcriptome analysis. Mirjam van Bentum: Generated and analyzed KO lines. Jasmine Hillmer: Performed data analysis. Jennifer von Schlichting: Performed data analysis. Reinhold Schäfer: Characterized cell lines and clones, manuscript revision. Nils Blüthgen: Supervision, funding acquisition, conceived the study, data analysis, manuscript writing. Michael Boettcher: Supervision, designed sgRNA library, designed experiments, funding acquisition, manuscript writing.

## SUPPLEMENTARY DATA

Supplementary Figures

Supplementary Data 1: Map of pMB1-10x vector for CRISPR screen sgRNA expression.

Supplementary Table 1: sgRNA Library sgRNA sequences

Supplementary Table 2: Proliferation screen read count table and MAGeCK MLE values

Supplementary Table 3: Modified TAP-seq targets inner primers for gene expression library amplification

Supplementary Table 4: sgRNA sequences used for the generation of EGR1 and FOS double knockout clonal lines

Supplementary Table 5: TCF7 signature genes identified from the Perturb-seq data

Supplementary Data are available at NAR online.

## DATA AVAILABILITY

Raw and processed transcriptome data is available at GEO under the accession number GSE250559. Data processing scripts and raw input data for the data processing scripts are available at Zenodo at the following doi: 10.5281/zenodo.10493549.

## REFERENCES

1. Schulze, A., Lehmann, K., Jefferies, H.B., McMahon, M. and Downward, J. (2001) Analysis of the transcriptional program induced by Raf in epithelial cells. Genes Dev., 15, 981–994.

2. Tullai, J.W., Schaffer, M.E., Mullenbrock, S., Sholder, G., Kasif, S. and Cooper, G.M. (2007) Immediate-early and delayed primary response genes are distinct in function and genomic architecture. J. Biol. Chem., 282, 23981–23995.

3. Tullai, J.W., Schaffer, M.E., Mullenbrock, S., Kasif, S. and Cooper, G.M. (2004) Identification of transcription factor binding sites upstream of human genes regulated by the phosphatidylinositol 3-kinase and MEK/ERK signaling pathways. J. Biol. Chem., 279, 20167–20177.

4. Jürchott, K., Kuban, R.-J., Krech, T., Blüthgen, N., Stein, U., Walther, W., Friese, C., Kiełbasa, S.M., Ungethüm, U., Lund, P., et al. (2010) Identification of Y-box binding protein 1 as a core regulator of MEK/ERK pathway-dependent gene signatures in colorectal cancer cells. PLoS Genet., 6, e1001231.

5. Amit, I., Citri, A., Shay, T., Lu, Y., Katz, M., Zhang, F., Tarcic, G., Siwak, D., Lahad, J., Jacob-Hirsch, J., et al. (2007) A module of negative feedback regulators defines growth factor signaling. Nat. Genet., 39, 503–512.

6. Avraham, R. and Yarden, Y. (2011) Feedback regulation of EGFR signalling: decision making by early and delayed loops. Nat. Rev. Mol. Cell Biol., 12, 104–117.

7. Legewie, S., Herzel, H., Westerhoff, H.V. and Blüthgen, N. (2008) Recurrent design patterns in the feedback regulation of the mammalian signalling network. Mol. Syst. Biol., 4, 190.

8. Uhlitz, F., Sieber, A., Wyler, E., Fritsche-Guenther, R., Meisig, J., Landthaler, M., Klinger, B. and Blüthgen, N. (2017) An immediate-late gene expression module decodes ERK signal duration. Mol. Syst. Biol., 13, 928.

9. Kim, E.K. and Choi, E.-J. (2010) Pathological roles of MAPK signaling pathways in human diseases. Biochim. Biophys. Acta, 1802, 396–405.

10. Dhillon, A.S., Hagan, S., Rath, O. and Kolch, W. (2007) MAP kinase signalling pathways in cancer. Oncogene, 26, 3279–3290.

11. Sanchez-Vega, F., Mina, M., Armenia, J., Chatila, W.K., Luna, A., La, K.C., Dimitriadoy, S., Liu, D.L., Kantheti, H.S., Saghafinia, S., et al. (2018) Oncogenic signaling pathways in the cancer genome atlas. Cell, 173, 321–337.e10.

12. Hibshman, P.S. and Der, C.J. (2024) The RAS signaling network and cancer. In Rauen, K.A. (ed), The rasopathies: genetic syndromes of the RAS/MAPK pathway. Springer Nature Switzerland, Cham, pp. 363–395.

13. Stelniec-Klotz, I., Legewie, S., Tchernitsa, O., Witzel, F., Klinger, B., Sers, C., Herzel, H., Blüthgen, N. and Schäfer, R. (2012) Reverse engineering a hierarchical regulatory network downstream of oncogenic KRAS. Mol. Syst. Biol., 8, 601.

14. Adamson, B., Norman, T.M., Jost, M., Cho, M.Y., Nuñez, J.K., Chen, Y., Villalta, J.E., Gilbert, L.A., Horlbeck, M.A., Hein, M.Y., et al. (2016) A Multiplexed Single-Cell CRISPR Screening Platform Enables Systematic Dissection of the Unfolded Protein Response. Cell, 167, 1867–1882.e21.

15. Dixit, A., Parnas, O., Li, B., Chen, J., Fulco, C.P., Jerby-Arnon, L., Marjanovic, N.D., Dionne, D., Burks, T., Raychowdhury, R., et al. (2016) Perturb-Seq: Dissecting Molecular Circuits with Scalable Single-Cell RNA Profiling of Pooled Genetic Screens. Cell, 167, 1853–1866.e17.

16. Schraivogel, D., Gschwind, A.R., Milbank, J.H., Leonce, D.R., Jakob, P., Mathur, L., Korbel, J.O., Merten, C.A., Velten, L. and Steinmetz, L.M. (2020) Targeted Perturb-seq enables genome-scale genetic screens in single cells. Nat. Methods, 17, 629–635.

17. El Kassem, G., Hillmer, J. and Boettcher, M. (2025) Evaluation of Cas13d as a tool for genetic interaction mapping. Nat. Commun., 16, 1631.

18. Replogle, J.M., Norman, T.M., Xu, A., Hussmann, J.A., Chen, J., Cogan, J.Z., Meer, E.J., Terry, J.M., Riordan, D.P., Srinivas, N., et al. (2020) Combinatorial single-cell CRISPR screens by direct guide RNA capture and targeted sequencing. Nat. Biotechnol., 38, 954–961.

19. Cagnol, S., Van Obberghen-Schilling, E. and Chambard, J.C. (2006) Prolonged activation of ERK1, 2 induces FADD-independent caspase 8 activation and cell death. Apoptosis, 11, 337–346.

20. Samuels, M.L., Weber, M.J., Bishop, J.M. and McMahon, M. (1993) Conditional transformation of cells and rapid activation of the mitogen-activated protein kinase cascade by an estradiol-dependent human raf-1 protein kinase. Mol. Cell. Biol., 13, 6241–6252.

21. McMahon, M. (2001) Steroid receptor fusion proteins for conditional activation of Raf-MEK-ERK signaling pathway. Meth. Enzymol., 332, 401–417.

22. Doench, J.G., Fusi, N., Sullender, M., Hegde, M., Vaimberg, E.W., Donovan, K.F., Smith, I., Tothova, Z., Wilen, C., Orchard, R., et al. (2016) Optimized sgRNA design to maximize activity and minimize off-target effects of CRISPR-Cas9. Nat. Biotechnol., 34, 184–191.

23. Sanson, K.R., Hanna, R.E., Hegde, M., Donovan, K.F., Strand, C., Sullender, M.E., Vaimberg, E.W., Goodale, A., Root, D.E., Piccioni, F., et al. (2018) Optimized libraries for CRISPR-Cas9 genetic screens with multiple modalities. Nat. Commun., 9, 5416.

24. Gibson, D.G., Young, L., Chuang, R.-Y., Venter, J.C., Hutchison, C.A. and Smith, H.O. (2009) Enzymatic assembly of DNA molecules up to several hundred kilobases. Nat. Methods, 6, 343–345.

25. Li, W., Xu, H., Xiao, T., Cong, L., Love, M.I., Zhang, F., Irizarry, R.A., Liu, J.S., Brown, M. and Liu, X.S. (2014) MAGeCK enables robust identification of essential genes from genome-scale CRISPR/Cas9 knockout screens. Genome Biol., 15, 554.

26. Hao, Y., Hao, S., Andersen-Nissen, E., Mauck, W.M., Zheng, S., Butler, A., Lee, M.J., Wilk, A.J., Darby, C., Zager, M., et al. (2021) Integrated analysis of multimodal single-cell data. Cell, 184, 3573–3587.

27. Ahlmann-Eltze, C. and Huber, W. (2021) glmGamPoi: fitting Gamma-Poisson generalized linear models on single cell count data. Bioinformatics, 36, 5701–5702.

28. Concordet, J.-P. and Haeussler, M. (2018) CRISPOR: intuitive guide selection for CRISPR/Cas9 genome editing experiments and screens. Nucleic Acids Res., 46, W242–W245.

29. Ran, F.A., Hsu, P.D., Wright, J., Agarwala, V., Scott, D.A. and Zhang, F. (2013) Genome engineering using the CRISPR-Cas9 system. Nat. Protoc., 8, 2281–2308.

30. Sell, T., Klotz, C., Fischer, M.M., Astaburuaga-García, R., Krug, S., Drost, J., Clevers, H., Sers, C., Morkel, M. and Blüthgen, N. (2023) Oncogenic signaling is coupled to colorectal cancer cell differentiation state. J. Cell Biol., 222.

31. Shaulian, E. and Karin, M. (2001) AP-1 in cell proliferation and survival. Oncogene, 20, 2390–2400.

32. Regan, J.L., Schumacher, D., Staudte, S., Steffen, A., Lesche, R., Toedling, J., Jourdan, T., Haybaeck, J., Golob-Schwarzl, N., Mumberg, D., et al. (2022) Identification of a neural development gene expression signature in colon cancer stem cells reveals a role for EGR2 in tumorigenesis. iScience, 25, 104498.

33. Kervestin, S. and Jacobson, A. (2012) NMD: a multifaceted response to premature translational termination. Nat. Rev. Mol. Cell Biol., 13, 700–712.

34. Cesana, M., Tufano, G., Panariello, F., Zampelli, N., Ambrosio, S., De Cegli, R., Mutarelli, M., Vaccaro, L., Ziller, M.J., Cacchiarelli, D., et al. (2023) EGR1 drives cell proliferation by directly stimulating TFEB transcription in response to starvation. PLoS Biol., 21, e3002034.

35. Margolin, A.A., Nemenman, I., Basso, K., Wiggins, C., Stolovitzky, G., Dalla Favera, R. and Califano, A. (2006) ARACNE: an algorithm for the reconstruction of gene regulatory networks in a mammalian cellular context. BMC Bioinformatics, 7 Suppl 1, S7.

36. Milo, R., Shen-Orr, S., Itzkovitz, S., Kashtan, N., Chklovskii, D. and Alon, U. (2002) Network motifs: simple building blocks of complex networks. Science, 298, 824–827.

37. Wang, B., Guo, H., Yu, H., Chen, Y., Xu, H. and Zhao, G. (2021) The role of the transcription factor EGR1 in cancer. Front. Oncol., 11, 642547.

38. Doumpas, N., Lampart, F., Robinson, M.D., Lentini, A., Nestor, C.E., Cantù, C. and Basler, K. (2019) TCF/LEF dependent and independent transcriptional regulation of Wnt/β-catenin target genes. EMBO J., 38.

39. Niida, A., Hiroko, T., Kasai, M., Furukawa, Y., Nakamura, Y., Suzuki, Y., Sugano, S. and Akiyama, T. (2004) DKK1, a negative regulator of Wnt signaling, is a target of the beta-catenin/TCF pathway. Oncogene, 23, 8520–8526.

40. Kim, D., Rath, O., Kolch, W. and Cho, K.H. (2007) A hidden oncogenic positive feedback loop caused by crosstalk between Wnt and ERK pathways. Oncogene, 26, 4571–4579.

41. Zeller, E., Hammer, K., Kirschnick, M. and Braeuning, A. (2013) Mechanisms of RAS/β-catenin interactions. Arch. Toxicol., 87, 611–632.

42. Zhan, T., Ambrosi, G., Wandmacher, A.M., Rauscher, B., Betge, J., Rindtorff, N., Häussler, R.S., Hinsenkamp, I., Bamberg, L., Hessling, B., et al. (2019) MEK inhibitors activate Wnt signalling and induce stem cell plasticity in colorectal cancer. Nat. Commun., 10, 2197.

