## Supplementary Figures for "Perturb-seq identifies TCF7 as a central nexus linking MAPK- and Wnt-driven gene expression"

### co-corresponding

#### Supplementary Figures

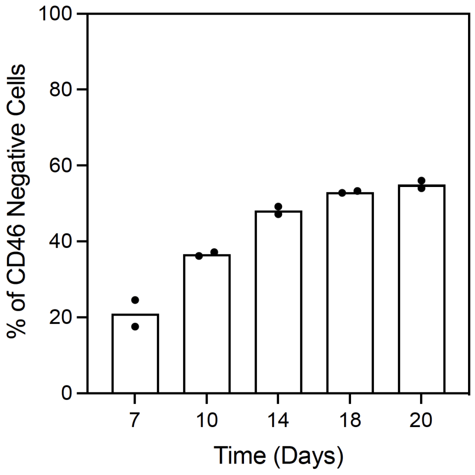


**Supplementary Figure 1: CD46 knockout kinetics in HEK293ΔRAF1:ER cells at different time points after lentiviral infection.** Percentage of CD46 knockout cells was determined via flow cytometry analysis of >10,000 cells stained with CD46 antibodies (Miltenyi).

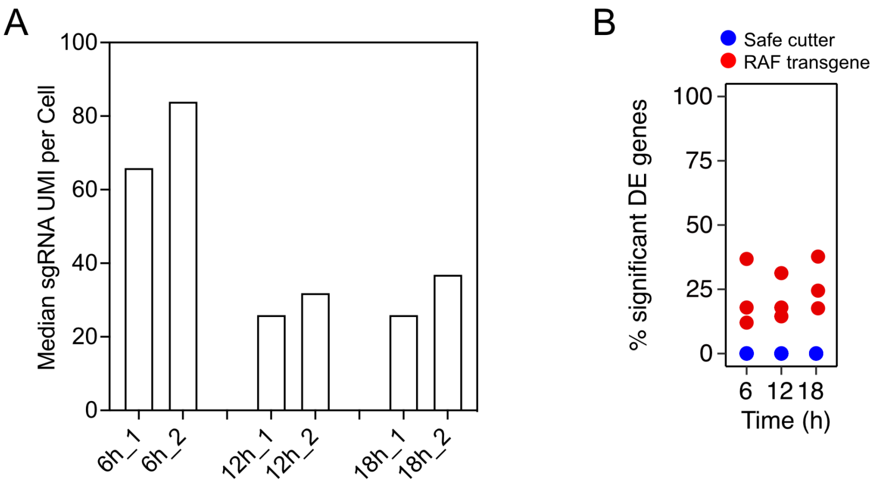


**Supplementary Figure 2: Quality assessment of Perturb-seq screen performance.** (A) Median sgRNA UMI counts per cell detected in the respective Perturb-seq samples. (B) Percentage of significantly differentially expressed genes in the RAF1-knockout cells from the total number of genes induced by RAF activation (adjusted p-value<0.05). Red = RAF1 sgRNAs, Blue = safe cutter sgRNAs.

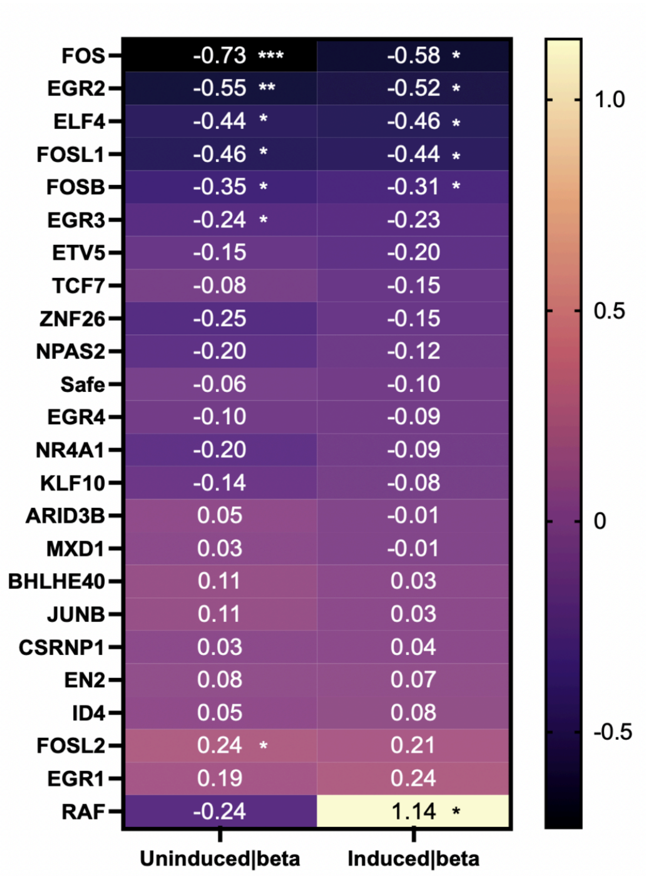


**Supplementary Figure 3: CRISPR/Cas9 proliferation screen in HEK293ΔRAF1:ER cells. MAGeCK MLE beta**

**scores and corresponding significance are shown.** * wald FDR <0.05, ** wald FDR <0.01, *** wald FDR <0.001

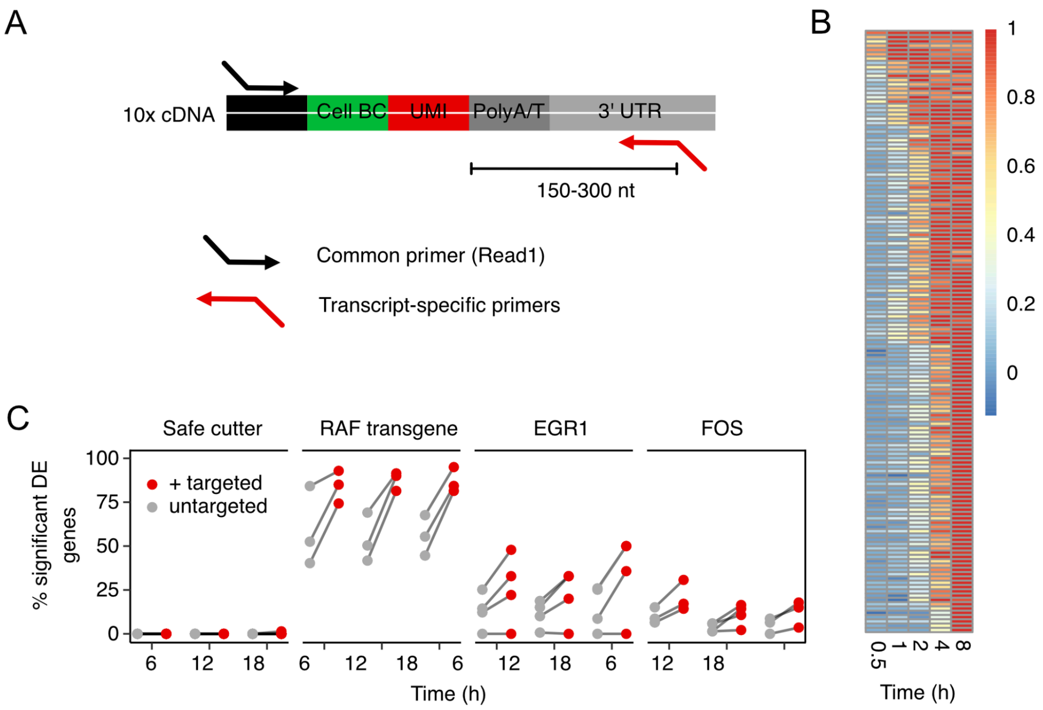


**Supplementary Figgure 4: Targeted scRNA-Seq library amplification enhances Perturb-seq sensitivity. (A)** Schematic of the modified TAP-seq approach. **(B)** Selection of 140 candidate genes for modified TAP-seq. $Log_2$ gene expression fold changes from bulk RNA-Seq analysis of significantly induced genes after pulse induction of the RAF transgene for different times (FDR = 1%), normalized to gene-wise maximum $Log_2$ fold changes. **(C)** Difference in percentage of significant differentially expressed genes from the 140 selected genes for the modified TAP-seq between targeted and untargeted approach after knockout of Raf transgene, EGR1, FOS, and safe cutter control.

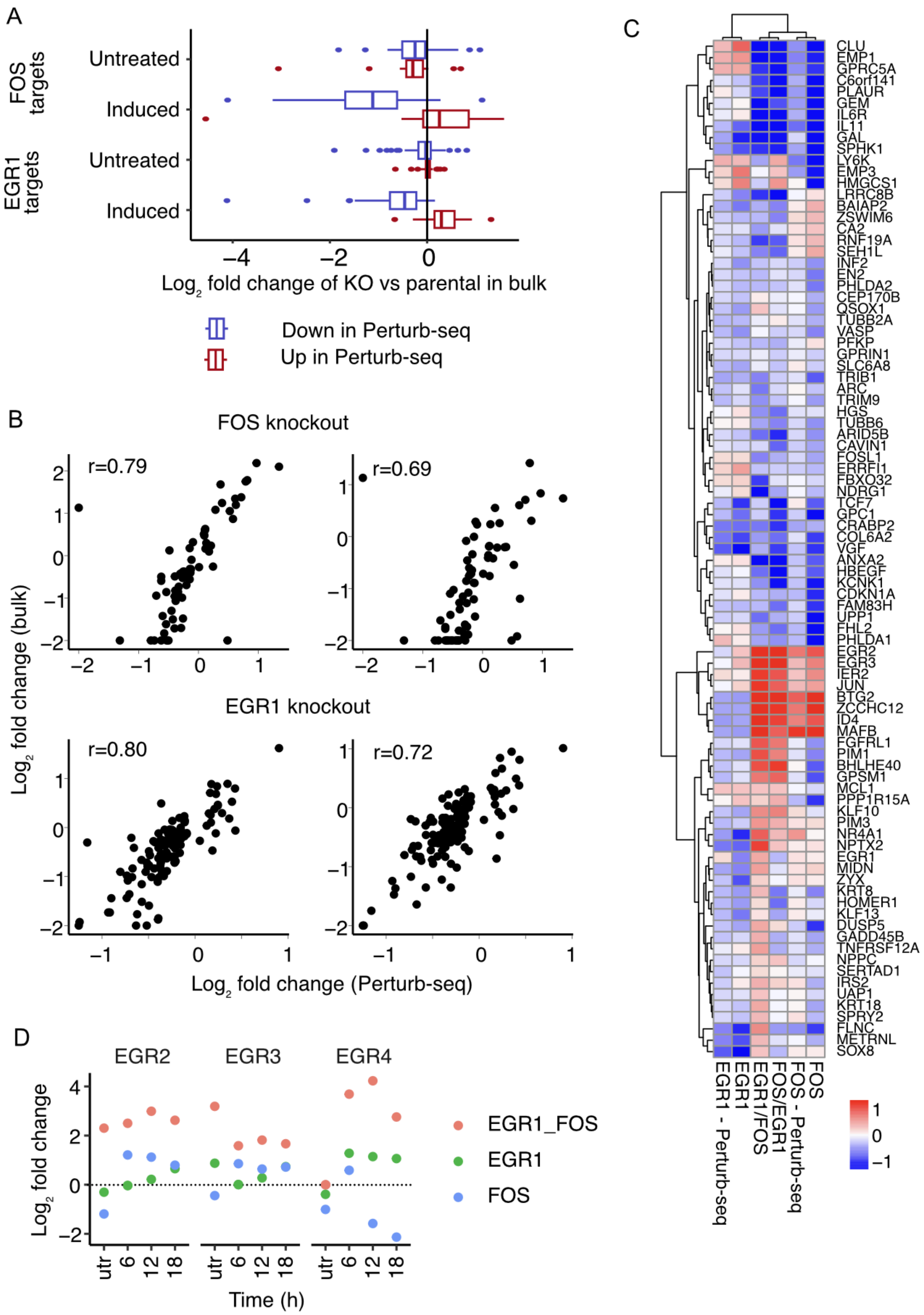

A
FOS targets
EGR1 targets
Untreated
Induced
Untreated
Induced
−4
−2
0
Log2 fold change of KO vs parental in bulk
Down in Perturb-seq
Up in Perturb-seq
B
FOS knockout
r=0.79
r=0.69
EGR1 knockout
r=0.80
r=0.72
Log2 fold change (bulk)
Log2 fold change (Perturb-seq)
C
CLU
EMP1
GPRC5A
C6orf141
PLAUR
GEM
IL6R
IL11
GAL
SPHK1
LY6K
EMP3
HMGCS1
LRRC8B
BAIAP2
ZSWIM6
CA2
RNF19A
SEH1L
INF2
EN2
PHLDA2
CEP170B
QSOX1
TUBB2A
VASP
PFKP
GPRIN1
SLC6A8
TRIB1
ARC
TRIM9
HGS
TUBB6
ARID5B
CAVIN1
FOSL1
ERRFI1
FBXO32
NDRG1
TCF7
GPC1
CRABP2
COL6A2
VGF
ANXA2
HBEGF
KCNK1
CDKN1A
FAM83H
UPP1
FHL2
PHLDA1
EGR2
EGR3
IER2
JUN
BTG2
ZCCHC12
ID4
MAFB
FGFRL1
PIM1
BHLHE40
GPSM1
MCL1
PPP1R15A
KLF10
PIM3
NR4A1
NPTX2
EGR1
MIDN
ZYX
KRT8
HOMER1
KLF13
DUSP5
GADD45B
TNFRSF12A
NPPC
SERTAD1
IRS2
UAP1
KRT18
SPRY2
FLNC
METRNL
SOX8
EGR1 - Perturb-seq
EGR1
EGR1/FOS
FOS/EGR1
FOS - Perturb-seq
FOS
1
0
−1
D
EGR2
EGR3
EGR4
Log2 fold change
EGR1_FOS
EGR1
FOS
utr
6
12
18
Time (h)

**Supplementary Figure 5: Bulk RNA-Seq analysis of individual and combinatorial EGR1 and FOS perturbations.** **(A)** Average $log_2$ fold changes for genes that are identified as up- and down-regulated target genes of EGR1 and FOS in perturb seq for bulk KO clones compared to parental clones. **(B)** Correlation between transcriptional changes detected via Perturb-seq and bulk RNA-Seq of EGR1- or FOS-perturbed cells from 2 different single knockout clonal lines of each gene. **(C)** Heatmap of transcriptional changes detected via bulk RNA-Seq from cells with individual or combinatorial EGR1 and FOS perturbations. Perturb-seq results from individual EGR1 and FOS perturbations are shown for comparison. **(D)** EGR2, EGR3, and EGR4 transcript levels determined via bulk RNA-Seq of EGR1/FOS single or double knockout clones, induced with 4OHT for 0h (utr), 6h, 12h, or 18h.
